# Two distinct chromatin modules regulate proinflammatory gene expression

**DOI:** 10.1101/2024.08.03.606159

**Authors:** Isabelle Seufert, Irene Gerosa, Vassiliki Varamogianni-Mamatsi, Anastasiya Vladimirova, Ezgi Sen, Stefanie Mantz, Anne Rademacher, Sabrina Schumacher, Panagiotis Liakopoulos, Petros Kolovos, Simon Anders, Jan-Philipp Mallm, Argyris Papantonis, Karsten Rippe

**Author notes:** Correspondence: Karsten Rippe.

## Abstract

Various mechanisms have been proposed to explain gene activation and co-regulation, including enhancer-promoter interactions via chromatin looping and the enrichment of transcription factors into hubs or condensates. However, these conclusions often stem from analyses of individual loci, and genome-wide studies exploring mechanistic differences with coupled gene expression are lacking. In this study, we dissected the proinflammatory gene expression program induced by TNFα in primary human endothelial cells using NGS- and imaging-based techniques. Our findings, enabled by our novel RWireX approach for single-cell ATAC-seq analysis, revealed two distinct regulatory chromatin modules: autonomous links of co-accessibility (ACs) between separated sites, and domains of contiguous co-accessibility (DCs) with increased local transcription factor binding. Genes in ACs and DCs exhibited different transcriptional bursting kinetics, highlighting the existence of two structurally and functionally distinct regulatory chromatin modules in the proinflammatory response. These findings provide a novel mechanistic framework for understanding how cells achieve rapid and precise gene expression control.

**Graphical abstract:** 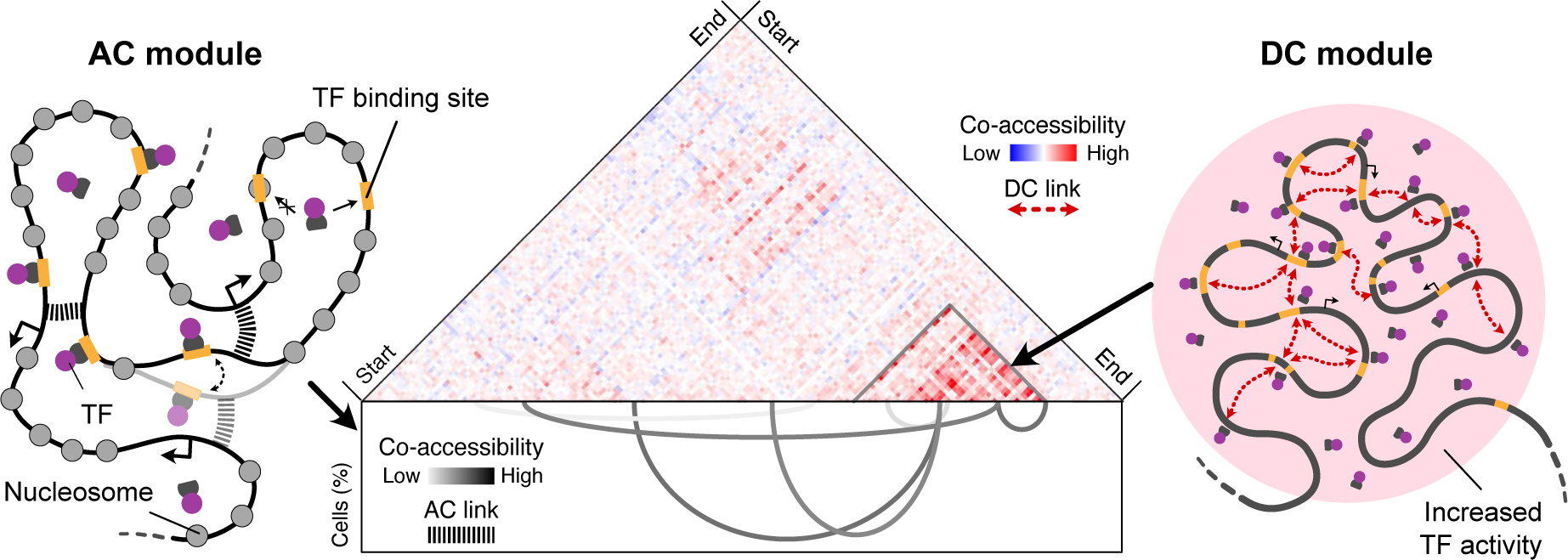

**Highlights:**

- Two distinct, non-mutually exclusive chromatin modules, ACs and DCs, that regulate proinflammatory gene expression were identified based on deep scATAC-seq.
- ACs represent long-range genomic interactions with regulation occurring more by transcription burst frequency.
- DCs are regions of increased local transcription factor binding that can modulate transcription burst size.
- The AC/DC model integrates sequencing-based evidence for chromatin looping with microscopy observations of transcription factor hubs/condensates into a unified model.
- Our findings provide a novel framework for understanding how cells achieve rapid and precise gene expression control.

## Introduction

Transcription in eukaryotes is a discontinuous process where genes alternate between bursts of activity and periods of silence ^1, 2^. These stochastic changes in gene activity can be jointly regulated across chromosomes through various mechanisms. Understanding the underlying mechanisms is crucial for deciphering how cells orchestrate rapid, precise, and coordinated responses to environmental stimuli, particularly those induced by inflammatory cytokines, where the timing and magnitude of gene expression are critical for cell viability.

Several models suggest that long-range interactions between cis-regulatory elements (CREs) such as promoters or enhancers ^3, 4^ play a crucial role in the clustering of RNA polymerase II (RNAP II) into transcription factories ^5^ or active chromatin hubs ^6^. The associated genome topology has been mapped by sequencing-based *in situ* cross-linking methods like Hi-C, which have identified topologically associating domains (TADs) as central structural units on the 0.1-1 MB scale ^7–12^. While these structural models provide insights into the spatial genome organization of transcriptional regulation, recent studies employing fluorescence microscopy and *in vitro* experiments have shifted the focus to the dynamic behavior and interactions of proteins involved in gene expression. These studies propose that phase separation of proteins and RNA drives the assembly of transcription factors (TFs), co-regulators, and the RNAP II machinery into protein assemblies termed “transcriptional condensates”, which accumulate at CREs and drive the transcriptional activity of multiple genes ^13–17^. Furthermore, single particle tracking experiments show that the chromatin microenvironment can confine TFs to specific regions of the nucleus, causing them to become locally enriched, pointing to the existence of “TF hubs” ^18–20^.

The above models are not mutually exclusive and may represent different aspects or scales of the same underlying regulatory mechanisms. However, a comprehensive, genome-wide analysis is still lacking that would provide a better understanding of how features from the various models could jointly contribute to direct gene expression programs. In particular, it remains unclear how different regulatory mechanisms, such as enhancer-promoter interactions, transcription factor dynamics, and local chromatin environments, work together to form “chromatin modules” as the functional units that direct complex transcriptional responses ^12, 21, 22^. Moreover, the relationship between different regulatory mechanisms and the observed patterns of transcriptional bursting is not well understood at a genome-wide level. Here, we employ a set of complementary single-cell sequencing readouts and fluorescence-based imaging to derive an integrated view of chromatin modules that drive the co-regulated induction of genes. We study the transcriptional response to tumor necrosis factor alpha (TNFα) treatment of human umbilical vein endothelial cells (HUVECs). The activation of the transcription factor NF-κB by TNFα induces a proinflammatory gene expression program, which represents a prototypical system for dissecting the linkage between gene regulation mechanisms and genome organization ^23–25^. We demonstrate how gene induction involves the co-regulation of genomically clustered genes, driven by two types of non-mutually exclusive chromatin modules: the “autonomous link of co-accessibility” (AC), which reflect a long-range interaction between separated CREs, and the “domain of contiguous co-accessibility” (DC), defined as a region of chromatin sites with increased local transcription factor activity rendered simultaneously accessible. Changes in transcriptional bursting kinetics upon TNFα treatment varied between genes in the AC versus DC modules, pointing to their functionally distinct regulatory mechanisms.

## Results

Transcription co-regulation was studied in HUVECs treated with TNFα for 0, 30, and 240 minutes, representing the uninduced, immediate-early, and later phases of the response. We performed single-cell/nuclei transcriptome analyses (scRNA-seq/snRNA-seq) and mapped open chromatin loci with the assay for transposase-accessible chromatin in nuclei (snATAC-seq) together with a multi-color single molecule FISH analysis of nascent transcripts. The single cell/nuclei analysis was complemented with bulk H3K27ac ChIP-seq data and the reanalysis of previously acquired 3’ bulk poly-A RNA-seq ^26, 27^ and Hi-C-seq data ^28^ (**Fig. 1A**). The 5’ scRNA-seq and snATAC-seq experiments were conducted in three independent triplicates at all three time points using our “TurboATAC” protocol for deep coverage of open chromatin sites ^29^. Cell cycle states were annotated to select cells in the G1 phase (**Fig. S1A-C**). This yielded homogeneous cell populations for each time point, which were used for all subsequent analyses and are visualized via UMAP embedding in **Fig. 1B**.

**Fig. 1.**
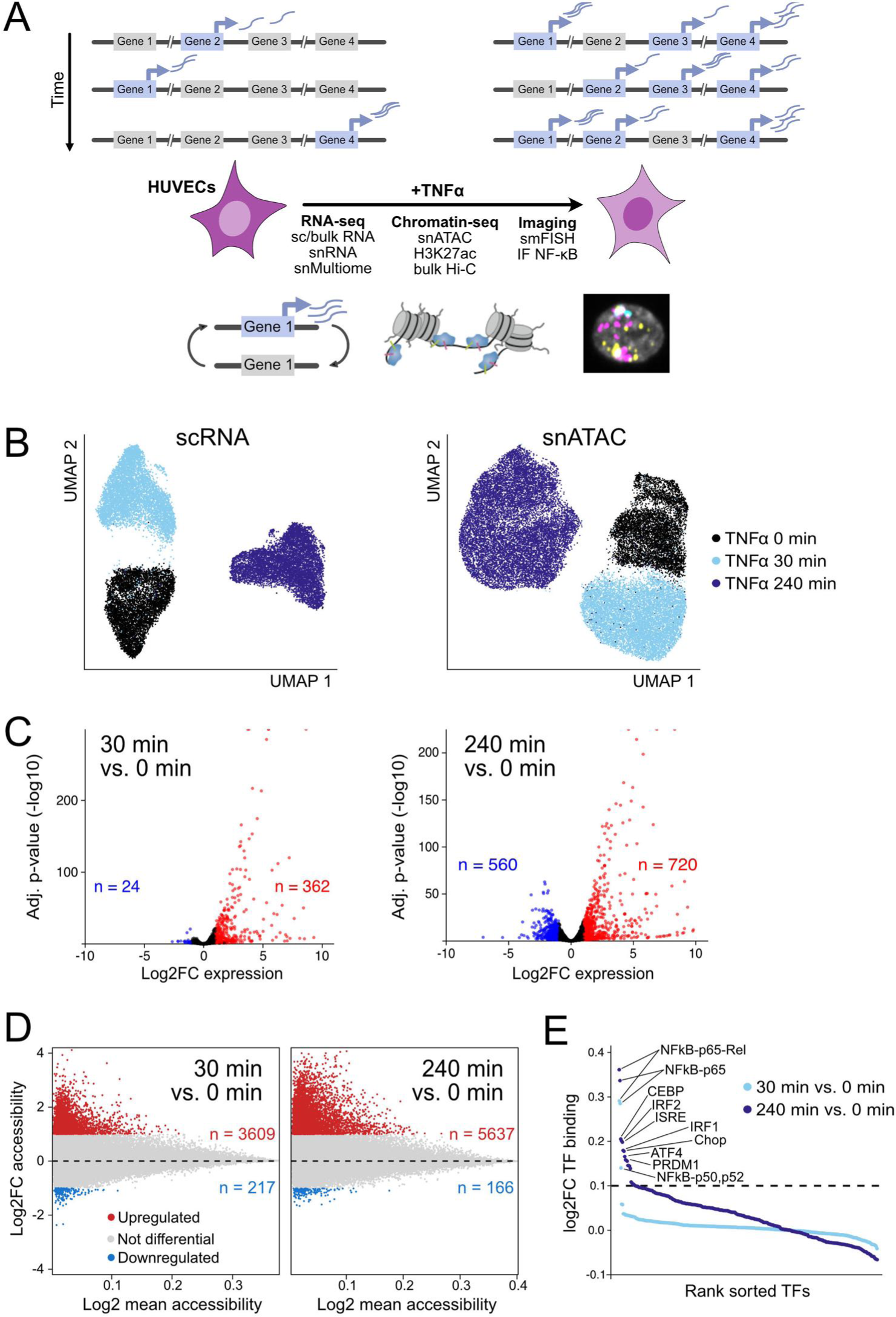
TNFα induced differences in gene expression and chromatin accessibility. (**A**) Changes in gene expression upon TNFα treatment were dissected in HUVECs with a complementary set of single cell and bulk sequencing readouts together with fluorescence microscopy of nascent RNAs and NF-κB. (**B**) UMAP embeddings of scRNA-seq (left) and snATAC-seq data (right) at 0 min (black), 30 min (light blue), and 240 min (dark blue) after TNFα induction. G1 cells from three biological replicates were selected to minimize the confounding effects of the cell cycle. (**C**) Gene expression changes upon TNFα induction across three biological replicates. Genes with log2 fold change (log2FC) ≥ 1 (up-regulated, red) and log2FC ≤ -1 (down-regulated, blue) and adjusted p-value < 0.05 were selected as TNFα regulated genes (TRGs). (**D**) Accessibility changes at ATAC peaks upon TNFα induction across three biological replicates. Peaks with differential accessibility of log2FC ≥ 1 (up-regulated, red) or log2FC ≤ -1 (down-regulated, blue) and adjusted p-value < 0.05 were used for further analysis. (**E**) Differential TF binding in ATAC peaks after TNFα treatment of HUVECs across three biological replicates. The dashed line separates TFs with binding log2FC <0.1, and the top 10 differential TFs are annotated.

### TNFα regulates ∼1,500 genes via NF-κB and IRF family TFs

Differential gene expression analysis was conducted using pseudo-bulks of scRNA-seq replicates for the 0-30 and 0-240 min time point comparisons (**Fig. 1C).** We identified 1,499 differentially expressed genes that are referred to as TRGs for TNFα-regulated genes (**Supplementary Dataset S2)**. This gene set included a significant fraction of down-regulated genes, which aligns with the observation that NF-κB, together with other TFs activated by TNFα, can act as an activator and a repressor, both directly and indirectly ^30, 31^. Our 5’ scRNA-seq data captured transcripts lacking poly(A) tails, identifying additional long non-coding RNAs (lncRNAs) as TRGs (∼30%). The heatmap of TRG expression across treatment conditions and biological replicates showed varying patterns of differential expression that distinguish between early and late (secondary) response kinetics (**Fig. S1D).** The expression analysis of 3’ bulk RNA-seq data identified fewer differentially expressed lncRNAs (3%) but confirmed differential expression of ∼70% of protein-coding TRGs (**Fig. S1E, F**). This comprehensive analysis reveals that TNFα regulates ∼1,500 genes, including a significant fraction of lncRNAs, highlighting its broad and complex impact on the inflammatory response.

### TNFα treatment induces the opening of NF-κB binding sites

Our snATAC-seq data analysis identified 201,329 chromatin accessibility peaks from a pseudo-bulk analysis that were located at promoters (9%), within gene bodies (61%), and in intergenic regions (30%) (**Fig. S1G**). At the single cell level, ∼150,000 unique fragments/cell were mapped, covering ∼50,000 peaks/cell with FRiP scores around ∼0.6 (**Supplementary Dataset S1**, **Fig. S1H**). Thus, our data provide deep open chromatin profiles of single cells, considering that only a fraction of the ∼200 thousand peaks from the aggregated data are simultaneously open in the same cell. The minus-average (MA) plots of differentially accessible peaks between 0-30 min and 0-240 min TNFα time points (**Fig. 1D**) revealed a relatively small fraction of 2-3% peaks with differential accessibility (3,826 peaks after 30 min; 5,803 peaks after 240 min). Of these, only 2% were at promoters, pointing to the importance of CREs located at intronic (∼55%) or intergenic sites (∼35%) (**Fig. S1G**). While 91% of all TRGs had an open chromatin site at the promoter, only 9% of these showed significant changes in promoter accessibility upon TNFα treatment (**Fig. S2A**). Pseudo-bulk analysis of differential TF binding revealed that NF-κB family motifs were strongly induced after 30 min of TNFα stimulation (**Fig. 1E**). In addition to NF-κB family motifs, after 240 min bona fide secondary targets of TNFα were induced, including IRF family, ATF4 (AP1), CEBP/CHOP and PRDM1 as the central motifs, consistent with previous studies on NF-κB crosstalk with other TFs ^32^. These findings demonstrate that TNFα induces targeted changes in chromatin accessibility, primarily at intronic and intergenic NF-κB binding sites, with only 2% of changes occurring at promoters, underscoring the critical role of distal regulatory elements in the inflammatory response.

### TRGs cluster in the genome and are co-induced

TRGs clustered along HUVEC chromosomes, which we visualized in a TRG network graph with proximal TRGs linked by edges (**Fig. 2A**). The number of TRG clusters varied with different distance thresholds to define local neighbors (**Fig. S2B**). At the 500 kb cutoff selected for further analysis, this resulted in 67% (1,008) TRGs in 356 clusters (**Supplementary Dataset S3**), while 33% (491) of TRGs were isolated. This number of TRG clusters exceeded the number expected for genomic clustering of randomly sampled genes (**Fig. S2C**). The TRG cluster average size was 460 kb, with most occupying less than 1 Mb (**Fig. S2D**). A cluster contained 2.8±1.3 TRGs on average, with a maximum of 9 TRGs (**Fig. S2D**). Most TRG clusters (69%) were located within a single TAD (**Fig. S2E**).

**Fig. 2.**
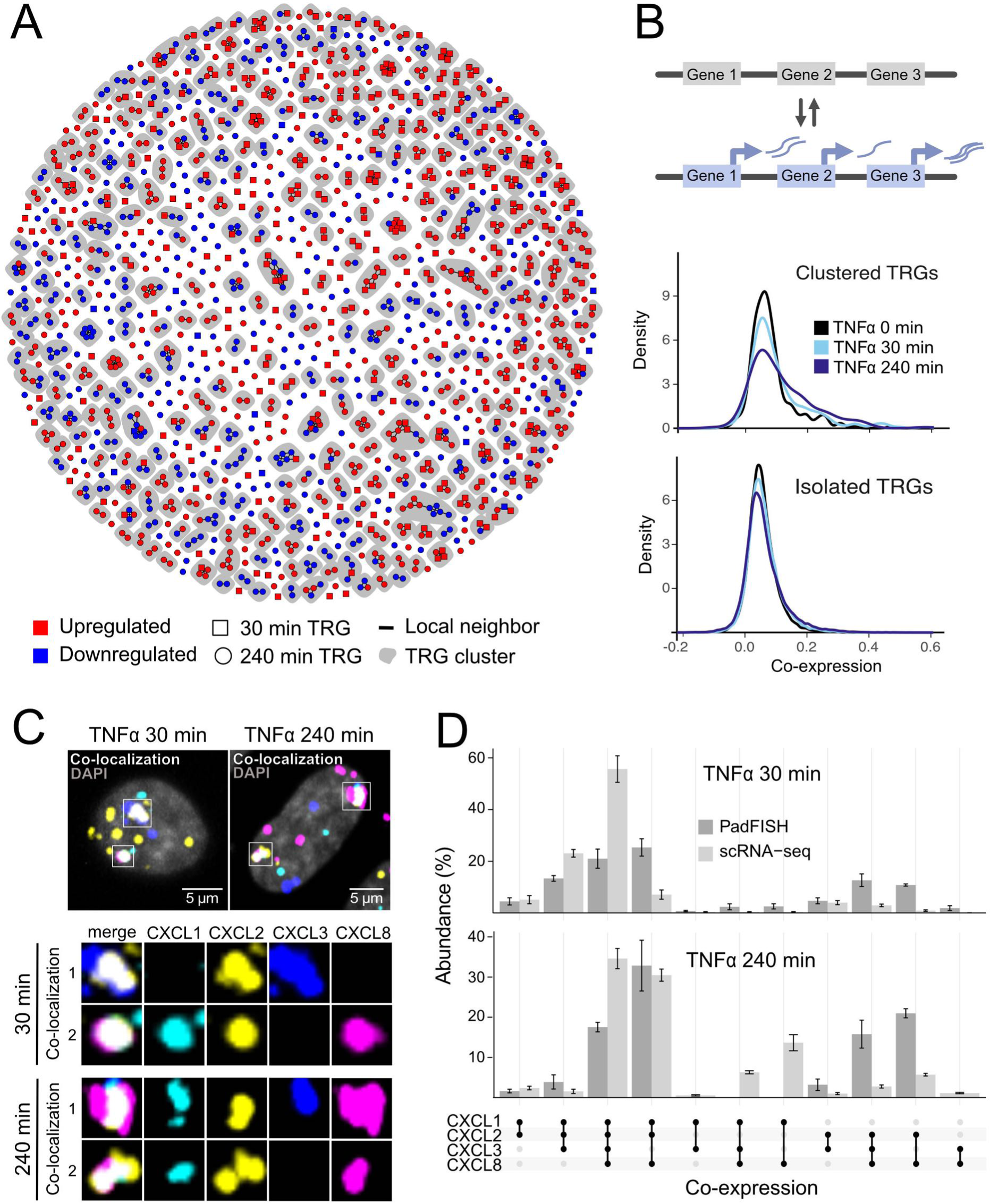
Genomic clustering and co-expression of TRGs. (**A**) Genomic clusters of TRGs below 500 kb distance. Each data point represents one TRG, with edges drawn to its neighbors. TRG color and shape indicate the direction and time point of differential expression. Shape and color are indicated for TRGs at both time points based on the 30 min results. Clusters of proximal TRGs are marked in grey. (**B**) Co-expression (replicate average) of clustered (top) and isolated (bottom) TRGs at the 0 min, 30 min, and 240 min time points. (**C**) padFISH images of intronic probes detecting nascent RNA from CXCL1 (cyan), CXCL2 (yellow), CXCL3 (blue), and CXCL8 (magenta) at 30 min and 240 min time points. Two exemplary cells are shown with a zoom-in of co-localized expression loci that appear in white color in the merged image. (**D**) CXCL co-expression patterns at 30- and 240-minute time points from padFISH and scRNA-seq. Error bars display standard errors from triplicates.

The density curves of co-expression from scRNA-seq revealed low co-expression of clustered and isolated TRGs in the absence of TNFα treatment (**Fig. 2B**). Upon TNFα stimulation, co-expression strongly increased within TRG clusters but not between isolated TRGs (**Figs 2B, S2F**). This finding was corroborated by a multiplexed single molecule FISH (smFISH) analysis using intronic padlock probes, termed padFISH, to detect nascent RNAs of the *CXCL* gene cluster ^33^ as an exemplary case for TNFα-induced gene co-expression (**Figs 2C, S2G, Supplementary Table S1**).

Both methods identified the same combination of genes in the *CXCL* cluster being co-expressed, namely either *CXCL1/2/3/8*, *CXCL1/2/3,* or *CXCL1/2/8*. In contrast, combinations that included *CXCL3* and *CXCL8* but not *CXCL2* were hardly ever detected (**Fig. 2D**). Our scRNA-seq and padFISH co-expression results were highly correlated (Spearman correlation coefficients of 0.83 at 30 min, and 0.79 at 240 min) (**Fig. S2H**). The genomic clustering (67% of TRGs in 356 clusters) and induced co-expression of TRGs upon TNFα stimulation suggest that spatial proximity and additional factors play a crucial role in facilitating coordinated gene regulation during inflammation.

### Co-accessibility analysis with RWireX reveals long-range features of gene regulation

Most TRGs displayed no differential chromatin accessibility at their promoters (**Fig. S1G, S2A**), suggesting that distal CREs are essential for the TNFα-regulated expression program. To gain further insight into the underlying mechanisms, we exploited the deep coverage of our snATAC-seq data. We developed the RWireX software package to map sites simultaneously accessible in the same cell as a proxy for regulatory interactions (**Fig. 3A, Supplementary methods**). A co-accessibility analysis with RWireX was conducted using two different workflows. The “single cell co-accessibility” workflow uses a homogeneous population of cells as input to identify co-accessible sites from stochastic accessibility changes between single cells. RWireX identified co-accessible links between high-resolution ATAC peaks by computing their correlation coefficients and percent accessible cells against a background model from shuffled input matrices.

**Fig. 3.**
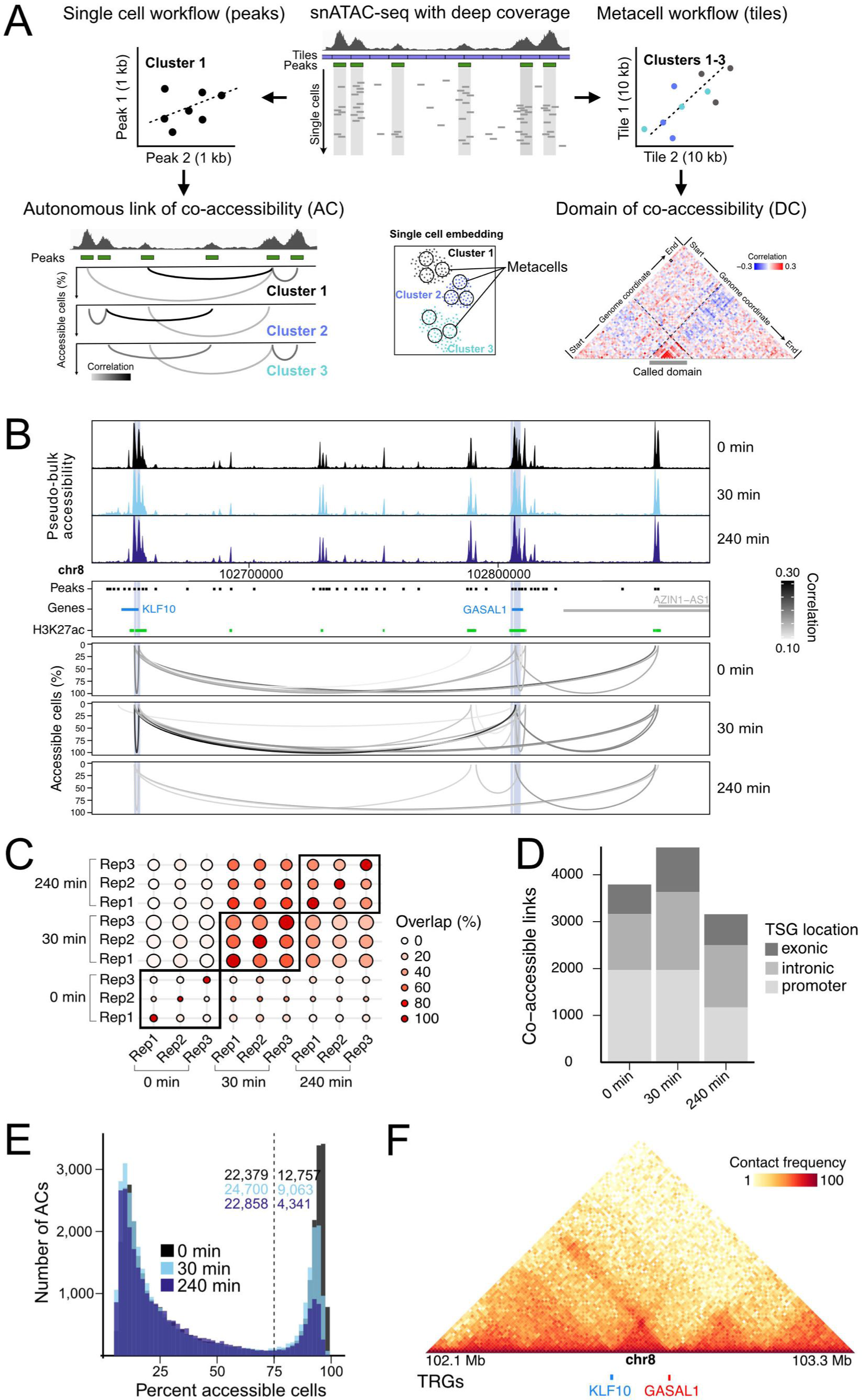
Co-accessibility analysis with RWireX. (**A**) RWireX computes co-accessibility from snATAC-seq data with two different workflows. Left: The single-cell co-accessibility is calculated between peaks. The grey levels visualize the magnitude of the corresponding correlation between two peaks. The height of the arcs indicates the percentage of cells in which at least one of the linked peaks is accessible. Right: The metacell co-accessibility is computed with aggregated cells in 10 kb genomic bins to identify domains with increased co-accessibility. (**B**) Single-cell co-accessibility at the KLF10/GASAL1 TRG cluster. Top: pseudo-bulk chromatin accessibility. Middle: pseudo-bulk ATAC peaks extended to 1 kb (black), H3K27ac peaks from ChIP-seq at 30 min time point (green), genes (grey), TRGs (blue), and 1 kb regions around their TSSs (light blue). Bottom: consensus autonomous links of co-accessibility (ACs) at TRG promoters visualized as described in panel A. (**C**) Replicates of ACs at ten most differential TRGs after 30 min of TNFα stimulation. The size and color of the dots show the total number of ACs detected in the reference sample and the percent overlap between the samples. (**D**) Number of consensus ACs at TRGs and their genomic location for TNFα time points. (**E**) Percent accessible cells at start and end peaks of ACs for the three different time points. (**F**) Chromatin contact map from HiC-seq data of unstimulated HUVECs. A region of 1.2 Mb around the KLF10/GASAL1 TRG cluster is shown with the upper color scale limit set to 100.

Additionally, we computed link activity scores for each cell, assessing whether both interacting peaks were detected as “open”. In contrast, the “metacell co-accessibility” workflow uses aggregated profiles of 10 cells with similar chromatin accessibility profiles to compute correlation coefficients between genomic tiles at a lower genomic resolution. For this, we analyzed heterogeneous cell populations with respect to a given perturbation, such as the duration of TNFα treatment. This approach allowed the identification of contiguous domains of co-accessibility enrichment in the genome driven by TNFα treatment, while any stochastic changes present in individual cells are no longer resolved. In this manner, the RWireX analysis uncovers both long-range regulatory interactions and larger regions of locally increased accessibility to dissect the chromatin-level mechanisms coordinating the TNFα-induced transcriptional response.

### TRG promoters and enhancers are frequently co-accessible in single cells

Single-cell co-accessibility analysis with RWireX was performed to detect long-range interactions between genomic loci. For example, a 160 kb region around the TRGs *KLF10* (log2FC_30 min_ = 3.04, log2FC_240 min_ not significant) and *GASAL1* (log2FC_30 min_ = 1.3, log2FC_240 min_ not significant) is shown in **Fig. 3B**. The two genes showed a co-accessible link between their promoters already before stimulation, which was present in almost all cells. This frequent co-accessible link increased markedly in strength after 30 min of TNFα treatment, and additional links to a potential intergenic enhancer, marked by H3K27ac enrichment, appeared. After 240 min of TNFα treatment, these links mostly disappeared, or their strengths again dropped, coinciding with the return of *KLF10* and *GASAL1* to basal expression levels.

The overlap of co-accessible links between replicates was high at TRGs (75%), but less so genome-wide (10%) (**Figs 3C, S3A, S3B**). Accordingly, we used links present in at least two replicates to compile consensus lists of autonomous links of co-accessibility (ACs) at each treatment time point for further analysis (**Fig. S3C**, **Supplementary Dataset S4**). We observed 12% of ACs at TRGs (10.8% at 0 min, 13.6% at 30 min, and 11.6% at 240 min; **Fig. S3D**), of which 45% were at the respective promoters, 36% within introns, and 19% in exons (**Fig. 3D**). The fraction of ACs between TRGs and distal H3K27ac sites was significantly higher (40%) than for non-TRG links (35%) (Chi-squared test p-value = 2.8e-11; **Fig. S3E**). The frequency at which ACs were detected in single cells displayed a well-separated bimodal distribution (**Fig. 3E**). A fraction of ACs was present in almost all cells, likely representing pre-established architectural interactions. In contrast, others showed a rare and more stochastic occurrence as they were detected in only a fraction of cells. Interestingly, the number of rare ACs remained constant throughout the treatment time course, while the number of frequent ACs decreased with ongoing TNFα treatment.

We then assessed the location of ACs in relation to TADs. While ACs were mainly located within the same TAD (45-52%), a significant fraction also extended across TAD boundaries (26-33%) or was found outside of TADs (23%) (**Fig. S3F**). Interestingly, the *KLF10* and *GASAL1* TRGs were located at the very boundaries of the same TAD (**Fig. 3F**). Finally, we investigated whether the presence of ACs correlated with gene expression by exploiting snMultiome-seq data (RNA and ATAC from the same nucleus), which were sparser than our separately acquired scRNA/snATAC-seq data. We computed Spearman correlation coefficients between TRG expression and their promoter’s link activities, as given by the multiplied accessibility of the two linked ATAC peaks (**Fig. S3G**). In general, the correlation between link activity and TRG expression was low. Nevertheless, the distribution displayed an extended right tail containing specific ACs that were correlated with TRG expression. Examples of links highly correlated with gene expression are shown for the key TRGs *JAG2*, *IER2* and *IRF1* (**Fig. S3H**). These findings, enabled by our novel RWireX approach for scATAC-seq analysis, reveal a complex landscape of promoter-enhancer interactions featuring both pre-established architectural links and more dynamic, stochastic connections.

### Metacell co-accessibility reveals domains of increased transcription factor activity

Next, we applied the RWireX metacell co-accessibility analysis and observed domains of contiguous co-accessibility (DCs) across the TNFα treatment time points. An example of a DC at the TRG cluster of *TNFAIP3*, *IFNGR1,* and lncRNA *WAKMAR2* is shown in **Fig. 4A**, referred to as the *TNFAIP3* DC in the following. All of these TRGs were significantly upregulated in response to TNFα (*TNFAIP3* log2FC_30 min_ = 5.9, log2FC_240 min_ = 4.8; *IFNGR1* log2FC_30 min_ not significant, log2FC_240 min_ = 1.4; *WAKMAR2* log2FC_30 min_ = 2.1, log2FC_240 min_ = 2.7) (**Fig. 4A**).

**Fig. 4.**
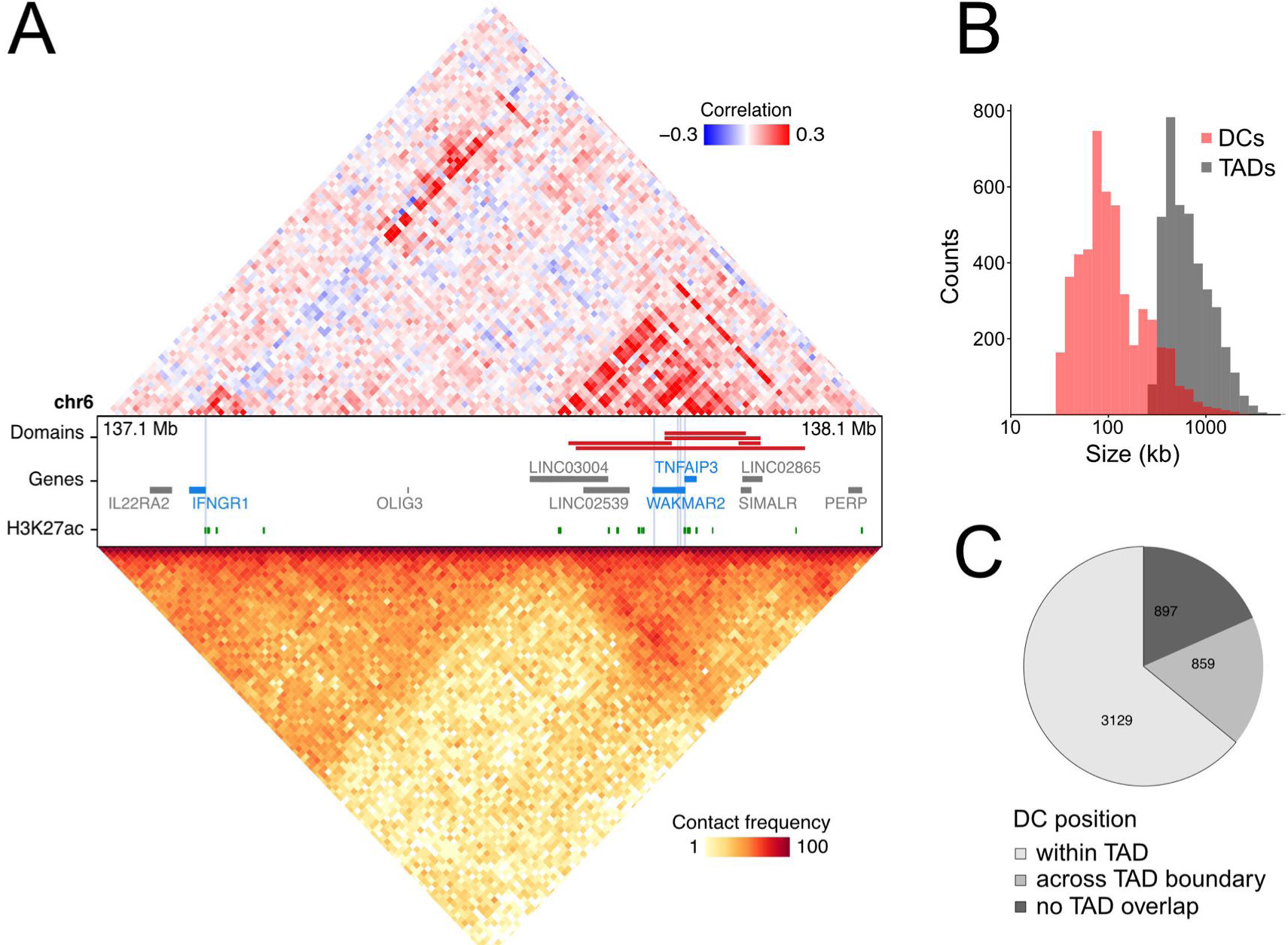
Co-accessibility analysis with RWireX using metacells. (**A**) Metacell co-accessibility (top) and chromatin contacts (bottom) at the TRG cluster of *TNFΑIP3*, *IFNGR1* and *WAKMAR2*. The metacell co-accessibility was computed across all time points, while the Hi-C data are from unstimulated HUVECs. The annotation in the middle shows DCs (black), H3K27ac peaks from ChIP-seq at 30 min time point (green), genes (grey), TRGs (blue) and 1 kb regions around their TSSs (light blue). (**B**) Genomic sizes of DCs and TADs. (**C**) Genomic location of DCs in relation to TADs. DCs were classified as within one TAD, across TAD boundary, and without TAD overlap.

Notably, TRG promoters in this cluster were devoid of ACs. Genome-wide, DCs were identified by repurposing the Hi-C TAD-calling tool SpectralTAD, which retrieved 4,885 domains based on the replicate consensus metacell co-accessibility (**Supplementary Dataset S5**).

The reproducibility of DC mapping in our replicate data was high, as illustrated by separate co-accessibility maps of each replicate in the exemplary region (**Fig. S4A**) and by >60% overlap between DCs from the individual replicates and the consensus (**Fig. S4B**). The size distributions of DCs and TADs indicated that DCs are TAD sub-structures (**Fig. 4B**). However, an analysis of the genomic location of DCs and TADs demonstrated that about 1/3 of DCs either overlapped with TAD boundaries or were located outside of TADs (**Fig. 4C**). Almost all DCs (95%) identified with RWireX showed significant accessibility changes upon TNFα treatment. The majority became less accessible upon TNFα treatment (74% at 30 min; 70% at 240 min), while only 26% (30 min) and 30% (240 min) showed increased accessibility in response to TNFα (**Fig. S4C**). However, this trend differed for the 683 DCs containing at least one TRG promoter. Three quarters of these displayed increased accessibility after 30 and/or 240 min of TNFα treatment, suggesting that TRG-containing DCs became activated. At the same time, the activity of DCs without TRGs was predominantly reduced. We then evaluated whether the presence of DCs directly correlated with gene expression in single cells. Spearman correlation coefficients between TRG expression and overall DC accessibility were computed, revealing a positive relationship between the two parameters (**Fig. S4D, S4E**).

Next, we investigated local TF binding activity in DCs using our pseudo-bulk snATAC-seq data (**Fig. 5A**). We computed TF footprints and inferred TF binding activity for the TFs that displayed a genome-wide increased binding activity upon TNFα treatment (**Fig. 1E**) using the TOBIAS software ^34^. We used TF binding activities to infer variations in TF binding in DCs against a local and a whole-genome non-DC background. Local enrichment of NF-κB binding was apparent when comparing the characteristic footprints of accessible NF-κB/p65 motifs within and around the merged *TNFAIP3* DC (**Fig. 5B**). Additionally, NF-κB/p65 binding activities of individual motifs in this DC were significantly higher than of the accessible motifs in the whole-genome non-DC background (**Fig. S5A-C**). Furthermore, a locally increased activity was observed for IRF family TFs, PRDM1, CEBP, and ATF4 (**Figs 5C, S5C**). Significant local enrichment for at least one differentially-bound TFs (**Fig. 1E**) was present in 44% of the DCs (**Fig. 5D**). Interestingly, the *TNFAIP3* DC showed high NF-κB binding activity already at the uninduced state. Immunostaining of NF-κB showed its targeting to the nucleus upon TNFα treatment and its assembly into nuclear foci where the protein is locally enriched (**Fig. S5C**). The nuclear concentration of NF-κB and the number of foci increased upon TNFα treatment (**Fig. S5D**). However, some nuclear NF-κB foci were apparent already at the uninduced time point. This observation aligns with NF-κB footprints at the 0 min time point from our sequencing-based analysis (**Figs S5E**), suggesting that a fraction of NF-κB DCs might be a persistent feature of HUVECs. Identifying DCs as local hubs of increased TF binding activity from scATAC-seq data unveils a novel layer in the spatial organization of transcriptional regulation during the inflammatory response.

**Fig. 5.**
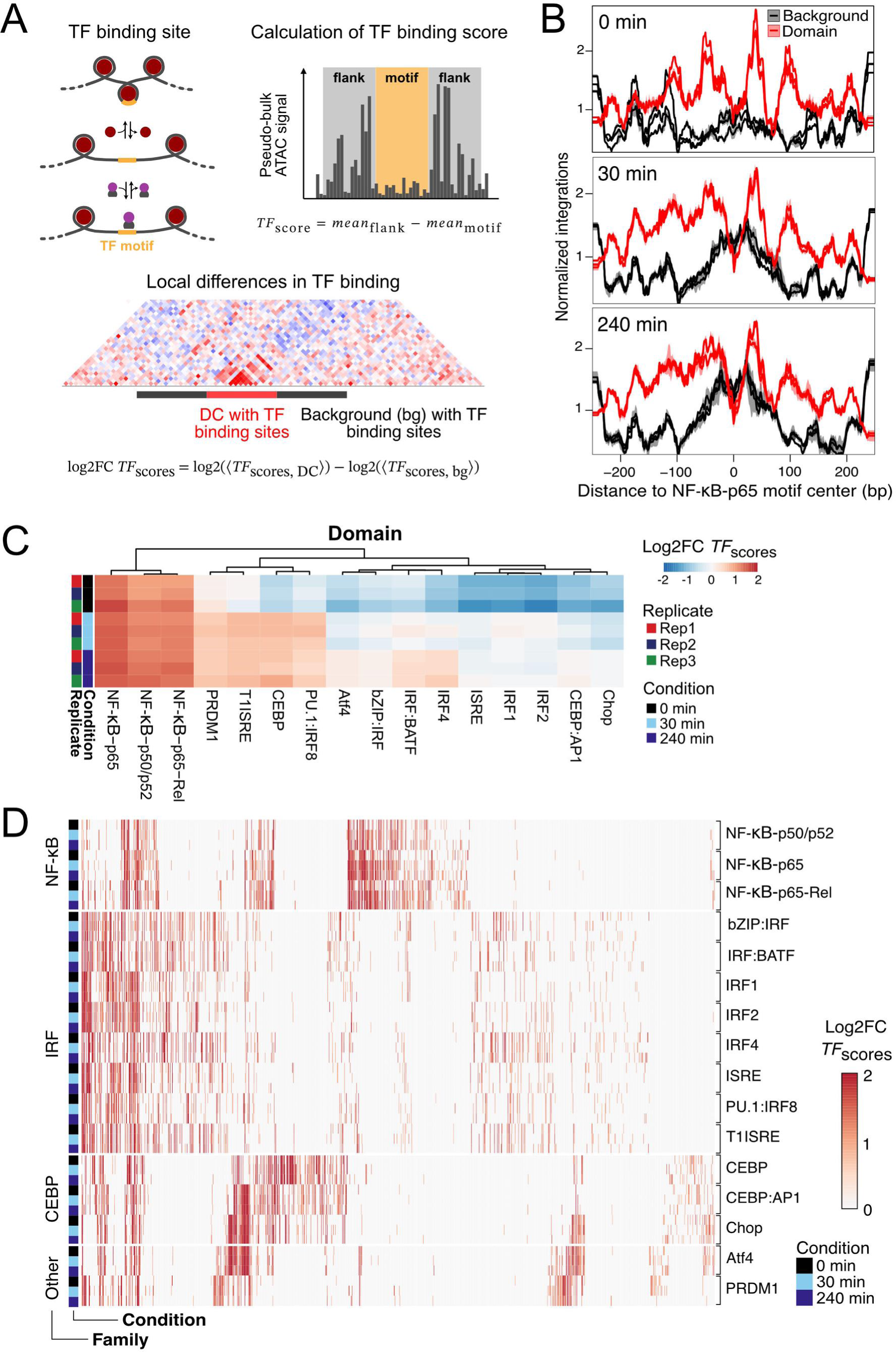
Local enrichment of TF binding activity in DCs. (**A**) Scheme of the approach to compute local TF binding activity in DCs. TF binding activity is inferred from pseudo-bulk footprints for each TF binding motifs in ATAC peaks. TF binding activity in DCs is compared to a local and whole-genome non-DC background. (**B**) Accessibility footprints at NF-κB/p65 motifs in the merged *TNFAIP3* DC (red) and the surrounding non-DC regions of the same size (local background; black). Each line shows the accessibility of one biological replicate in unstimulated (top), 30 min (middle), and 240 min (bottom) TNFα-simulated HUVECs. (**C**) Differential TF binding activity in the *TNFAIP*3 DC vs. the whole-genome non-DC background. Color scale limits are set to -2 and 2. (**D**) Same as panel C showing all DCs with significant local enrichment of TF binding activity from meta-analysis of replicates. TFs are grouped by family. DCs are clustered by summed family enrichment. The color scale limits are set to 0 and 2.

### Two distinct chromatin module types regulate TRG clusters

Based on our RWireX analysis, we annotated all TRGs with respect to their promoters having ACs and/or being in a DC. Subsequent clustering of TRGs distinguished four main groups, as visualized in **Fig. 6A**: DC-driven TRGs, AC-driven TRGs, TRGs with both AC/DC features, and TRGs carrying neither AC nor DC features. A comparison of the different regulation types showed increased DC regulation of clustered, upregulated, and early-response TRGs (**Fig. S6A, Datasets Supplementary Table S2**). In contrast, protein-coding, downregulated, and late-response TRGs displayed a preference for regulation via ACs. The analysis of nascent transcripts detected in purified transcription factories ^35^ displayed no apparent enrichment with respect to the AC or DC annotation. The *SAMD4* and *EXT1* genes previously associated with “NF-κB transcription factories” ^23^ were both in the AC/DC category. Next, we annotated TRGs in clusters identified above (**Fig. 2A**) concerning their regulation type as derived from the cluster composition (**Fig. 6B**).

**Fig. 6.**
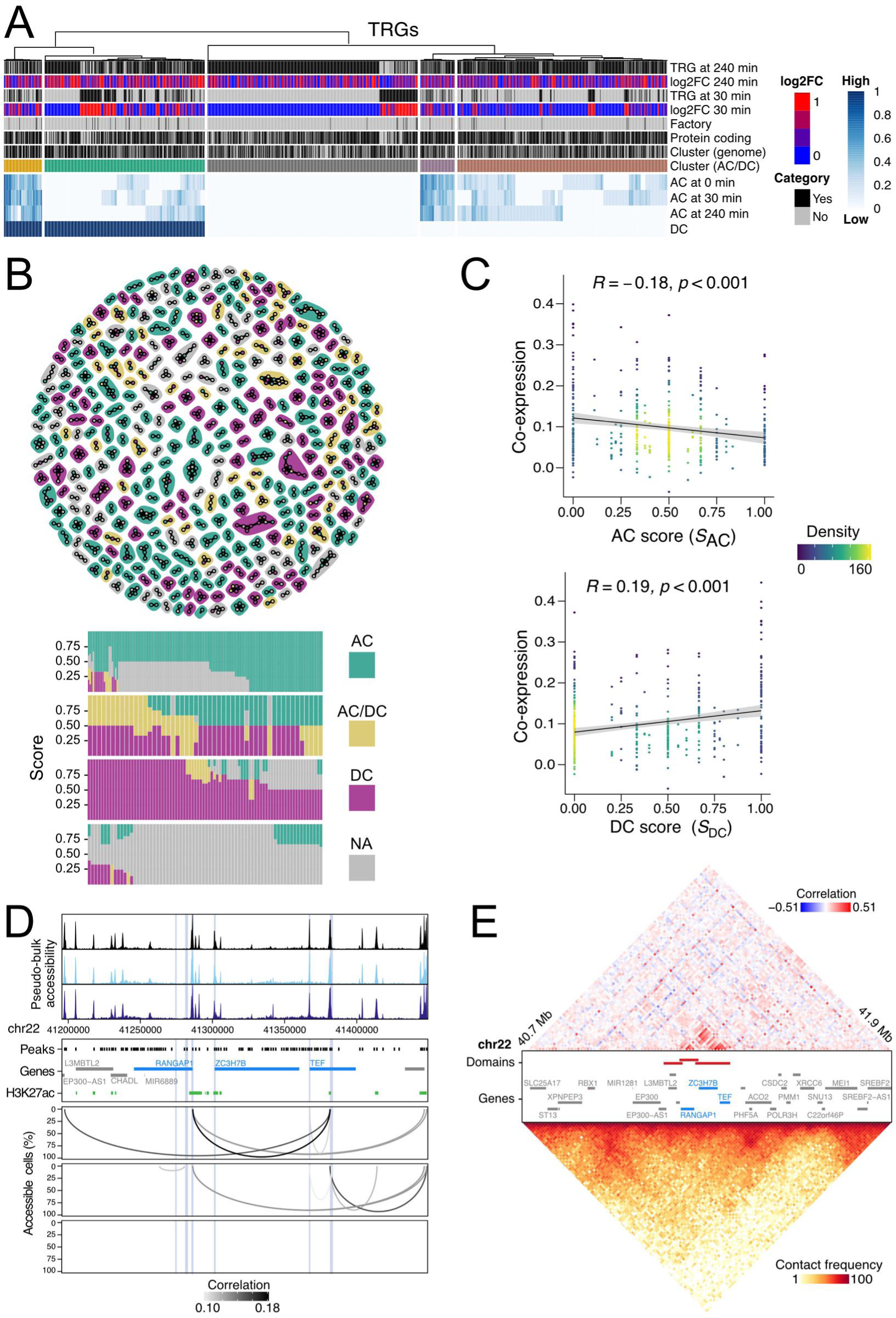
Chromatin modules at TRG clusters. (**A**) Clustering of TRGs into AC-, DC- and AC/DC-driven genes. The TRG annotation includes differential expression after 30 and 240 min of TNFα treatment, location (isolated/clustered), gene type (protein-coding/lncRNA), and detection of transcripts in purified transcription factories (factory). (**B**) TRG cluster types. Colors indicate the type of each TRG and the prevalent module type in the cluster. Top: network graph with each datapoint representing one TRG and edges to their local neighbors below 500 kb distance. Bottom: Composition of TRG clusters assigned to AC, DC, AC/DC, or NA (not assigned) chromatin modules. Each column represents one TRG cluster. (**C**) Co-expression in TRG clusters (replicate average) in dependence of their AC or DC scores (*S*_AC_, top; *S*_DC_, bottom). Colors reflect the density of TRG clusters. (**D**) Single-cell co-accessibility at the TRG cluster with ZC3H7B, RANGAP1, and TEF. Top: pseudo-bulk chromatin accessibility. Middle: pseudo-bulk ATAC peaks extended to 1 kb (black), H3K27ac peaks from ChIP-seq at 30 min time point (green), genes (grey), TRGs (blue), and 1 kb regions around their TSSs (light blue). Bottom: consensus ACs at TRG promoters. (**E**) Metacell co-accessibility (top) and chromatin contacts (bottom) at the TRG cluster with ZC3H7B, RANGAP1, and TEF. The metacell co-accessibility was computed across all time points, while the Hi-C data are from unstimulated HUVECs. The annotation in the middle shows DCs (black), genes (grey), TRGs (blue), and 1 kb regions around their TSSs (light blue).

An AC- and a DC-score (*S*_AC_ or *S*_DC_) was calculated for each TRG cluster. Based on these scores, TRG clusters were annotated as predominantly AC- or DC-driven or involving a combination of both (AC/DC) (**Fig. S6B**). TRGs within the same cluster were enriched for the same module type. AC, DC, and AC/DC modules displayed no significant differences in cluster size, TRG number, and TRG neighbors (**Fig. S6C**). A scatter plot of TRG cluster co-expression versus the *S*_DC_ DC-score showed a positive correlation coefficient of 0.19 (**Fig. 6C**). Thus, TRG co-regulation via local TF enrichment increased co-expression. In contrast, a negative correlation of -0.18 was observed for TRG cluster co-expression and *S*_AC_-values, suggesting that AC chromatin modules do not promote gene co-expression. These differences could be related to the bimodal distribution of TRG cluster co-expression shown in **Fig. S2F** and point to a functional difference between the two chromatin modules.

Examples of these different regulatory architectures are given in **Figs 3B** (AC) and **4A** (DC). In addition, a 150 kb region around the late-responsive TRGs *ZC3H7B* (log2FC_240 min_ = 2.2), *RANGAP1* (log2FC_240 min_ = 1.7), and *TEF* (log2FC_240 min_ = 1.4) is shown in **Figs 6D and 6E**, as an example for an AC/DC chromatin module. Interestingly, the ACs present at 0 and 30 min were lost after 240 min of TNFα treatment (**Fig. 6D**), suggesting that they could be associated with a repressive chromatin state. At the same time, a DC comprising the TRG promoters overlapped with a region of increased Hi-C contacts lacking clear boundaries (**Fig. 6E**). Thus, our analysis distinguishes the AC and DC type of regulatory chromatin modules in TRG clusters that can also co-exist at the same cluster (AC/DC). These findings provide a novel mechanistic framework for understanding how cells achieve rapid and precise gene expression control.

### AC and DC modules correlate with distinct 3D chromatin organization features

Next, we investigated the relation of ACs and Hi-C contacts in further detail. We computed the density curves of all Hi-C contacts and Hi-C contacts at ACs (**Fig. 7A**). This revealed a bimodal distribution with a fraction of ACs that had Hi-C contact frequencies ∼50 times than the genome-wide average. These sites corresponded to ACs within TADs or not within TADs. At the same time, Hi-C contact frequencies were largely reduced for ACs across TAD boundaries (**Fig. 7B**). Additionally, we investigated Hi-C contacts at TADs and the DC location. In aggregate peak analysis plots ^28^, Hi-C contacts of scaled TADs were averaged to compare TADs with DCs within and across their boundaries (**Fig. 7C**) and all TADs without DCs (**Fig. S7A**). In this analysis, TAD boundaries appeared weaker when overlapping with DCs, and increased interactions with neighboring TADs were observed.

**Fig. 7.**
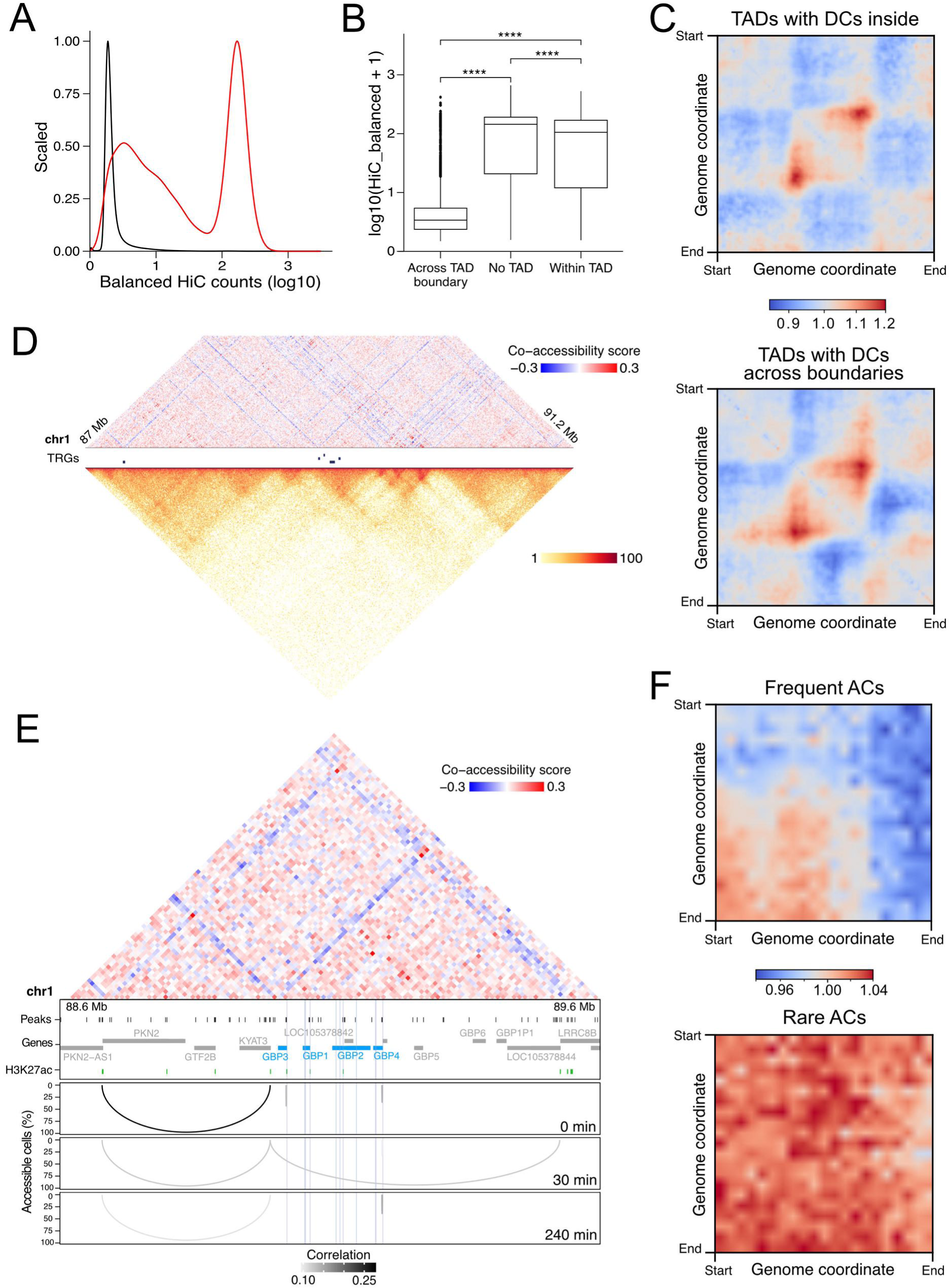
Hi-C chromatin contacts in unstimulated HUVECs at AC and DC chromatin modules. (**A**) Chromatin contacts genome-wide (black) and between AC-linked peaks (red). (**B**) Chromatin contacts between AC-linked peaks within TADs, across TAD boundaries, and outside TADs. P-values < 2.22e-16 from Wilcoxon test are indicated by ****. (**C**) Chromatin contact pileups of TADs with DCs within (top) and across TAD boundaries (bottom). (**D**) Metacell co-accessibility (top) and chromatin contacts (bottom) at the GBP TRG cluster. TRGs are annotated in blue. (**E**) Zoom in on the GBP cluster (central region from panel D) with metacell co-accessibility (top) and single-cell co-accessibility (bottom). The annotation in the middle shows pseudo-bulk ATAC peaks (black, extended to 1 kb), H3K27ac peaks from ChIP-seq after 30 min TNFα treatment (green), genes (grey), TRGs (blue), and 1 kb regions around their TSSs (light blue). (**F**) Chromatin contact pileups of frequent (top) and rare (bottom) ACs.

A metacell co-accessibility map of the *GBP* TRG cluster displayed lines of anti-correlated accessibility (blue color) with high co-accessibility at their junctions (**Fig. 7D**). These ‘blue borders’ often originated from genomic loci with gene promoters in the proximity of TAD boundaries. They coincided with stripes of increased chromatin interactions in Hi-C data. Zooming into a smaller region of this metacell co-accessibility map with annotated H3K27ac and ACs in the *GBP* cluster (**Fig. 7E**) showed that the blue borders represent distinct ACs between sites enriched for H3K27ac that were present in nearly 100% of cells. As an additional example, H3K27ac peaks and gene promoters were also located at the origins of such blue borders in the metacell co-accessibility map for the *KLF4* TRG locus (**Fig. S7B, S7C**). Again, the blue borders coincided with ACs present in nearly 100% of cells, and Hi-C contact maps further confirmed their coincidence with stripes of increased chromatin interactions. These findings suggest that the blue borders are linked to the frequently occurring and potentially architectural ACs described above (**Fig. 3E**) and might represent a subset of AC chromatin modules (*GBP1/3/4* are AC; *GBP2* is NA; *KLF4* is AC). To assess their spatial chromatin interactions, aggregate peak analysis plots of Hi-C contacts at AC interactions were scaled and averaged to compare Hi-C contacts at rare and frequent ACs (**Fig. 7F**). Rare ACs showed uniform Hi-C contacts in their entire vicinity while frequently occurring ACs showed distinct enrichment of Hi-C contacts between the linked sites. Overall, such blue borders could reflect stacking of loops/TAD boundaries ^36, 37^, but their specific underlying spatial relations that lead to the distinct co-accessibility pattern observed here cannot be inferred from this data. These observations reveal distinct relationships between ACs, DCs, and Hi-C chromatin contacts that provide insights into how different gene regulatory modules may leverage 3D genome organization: A fraction of ACs located in TADs displayed a largely increased Hi-C contact frequency, DCs spanning two TADs are associated with weakened TAD boundaries, and frequently occurring ACs correlate with an increase in Hi-C contacts between the linked sites.

### AC and DC chromatin modules differ in transcriptional bursting response to TNFα

Last, we tested whether the location of a TRG in AC, DC or AC/DC chromatin modules was related to its bursting kinetics. A two-state model of transcriptional bursting was applied that yielded the burst frequency rate *k*_on_ and the burst size from the *k*_syn_/*k*_off_ ratio according to the mechanism depicted in **Fig. 8A**. These parameters were first computed for each time point from intronic snRNA-seq reads. Scatter plots and density distributions of bursting kinetics revealed higher burst frequencies of TRGs in AC and AC/DC chromatin modules than TRGs in DC chromatin modules at all time points (**Figs 8A, S8A**). In contrast, the burst sizes in DC and AC/DC modules were higher than in AC modules after 30 and 240 min of TNFα treatment (**Figs 8A, S8A**). In line with these differences, the log_2_FC values of each TRG after 30 or 240 min of TNFα treatment showed a predominant regulation of DC TRGs by burst size (**Fig. S8B**).

**Fig. 8.**
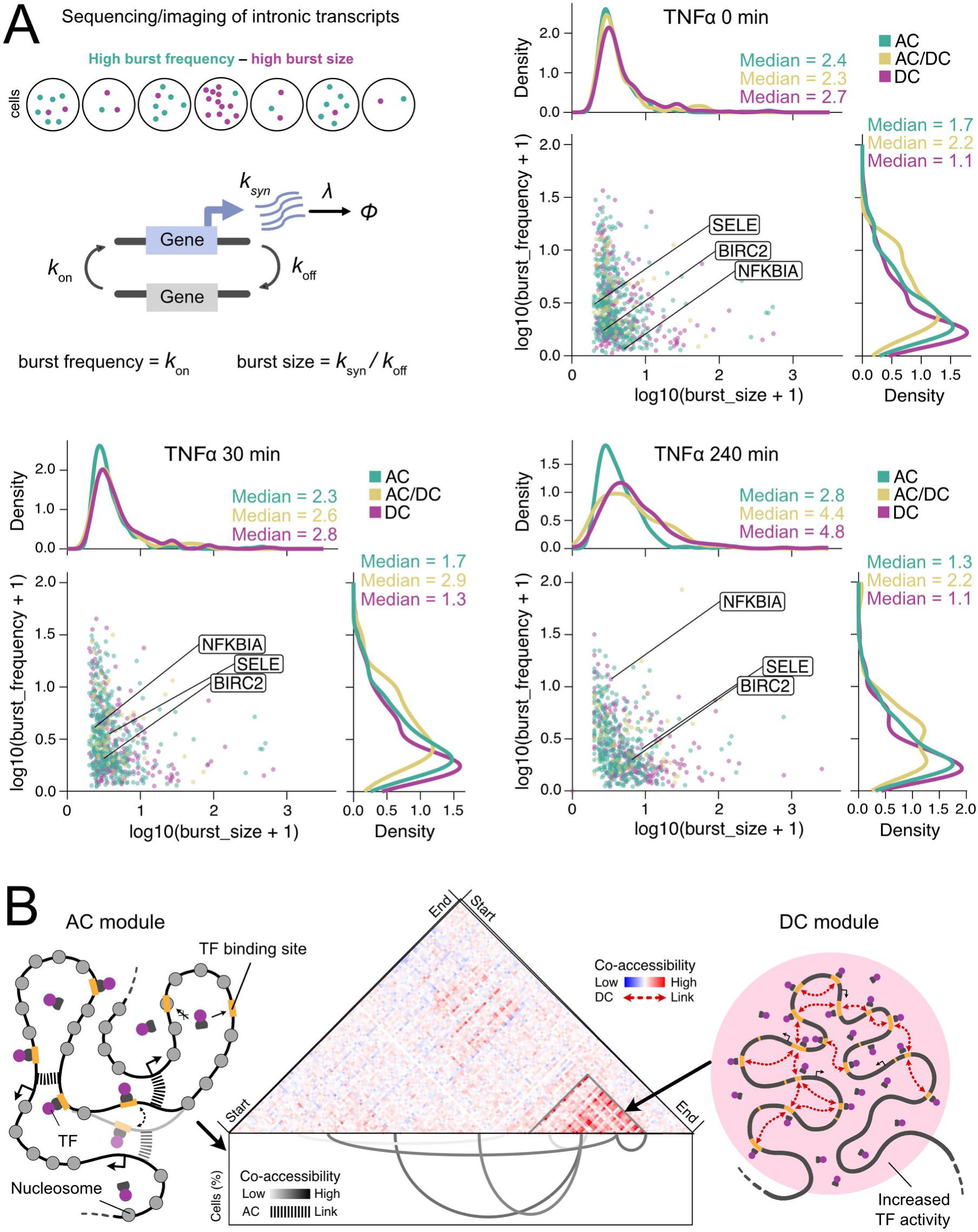
Functional differences in transcriptional burst kinetics of AC and DC modules. (**A**) Transcriptional bursting analysis according to the depicted model. TRG burst frequency and burst size (both log10 with pseudo-count of 1) at 0 min (top), 30 min (bottom left), and 240 min (bottom right) of TNFα treatment are shown. The colors in the scatter and density plots reflect the TRG’s chromatin module type. NFKBIA, SELE, and BIRC2 are highlighted as exemplary TRGs with further data provided in **Fig. S8**. **(B**) The AC/DC model of transcription regulation at TRG clusters. ACs are characterized by the enrichment of distinct co-accessibility correlations between 1 kb size ATAC peaks over larger distances. They occur in a stochastic manner but form two groups with low and high frequency of occurrence in cells. AC modules regulate transcription more frequently by an increased burst frequency, and TRGs in the same cluster show anti-correlated expression. DCs are domains of contiguous co-accessibility computed at 10 kb resolution that display an increased TF binding activity. Their presence correlates with TRG co-expression via changes in burst size.

We then compared the bursting kinetics of exemplary TRGs *NFKBIA*, *SELE*, and *BIRC2* derived from the snRNA-seq data to those inferred from single molecule FISH implemented via our padFISH protocol. Both methods yielded essentially the same results. *NFKBIA* showed low variation in burst size but a substantial increase in frequency across time points (**Fig. S8C, S8D**); *SELE* displayed an increase in burst size in padFISH and snRNA-seq, while burst frequencies remained stable (**Fig. S8E, S8F**); *BIRC2* showed an increase in burst size after 30 min in padFISH and after 30 and 240 min in snRNA-seq, but displayed hardly any changes in burst frequency (**Fig. S8G, S8H**). Thus, the padFISH analysis of *NFKBIA*, *SELE*, and *BIRC2* validates the approach of retrieving burst frequency and size from snRNA-seq data. We conclude that AC and DC modules employ functionally different transcription induction mechanisms, with ACs primarily affecting burst frequency and DCs influencing burst size.

### The AC/DC analysis is applicable to other cellular systems and perturbances

The approach introduced here to distinguish transcription regulation via AC and/or DC chromatin modules is generally applicable to snATAC-seq data with sufficiently deep coverage as illustrated in **Fig. S9** for three examples from different cellular systems using data from refs ^38, 39^. First, a DC identified at an interferon-induced gene cluster in mouse embryonic stem cells is shown, which contains the *Irf9*, *Psme1,* and *Psme2* genes (**Fig. S9A**). Second, the *Rnf213* gene induced by IFNβ stimulation and located in a DC in epithelial-like mouse embryonic fibroblasts is depicted (**Fig. S9B**). Third, a DC at the *Gimap6* gene (**Fig. S9C**), as well as frequently occurring ACs at the *Slc11a1* gene (**Fig. S9D**), were identified in a TCL1 mouse model upon *Tbx21* knockout leading to the loss of the T-bet transcription factor. These findings demonstrate that AC and DC modules can also be distinguished in other cellular systems that have different perturbances of their gene expression program.

## Discussion

Previous studies have shown that activation of the NF-κB pathway by TNFα or other cytokines is linked to chromatin reorganization underlying gene co-expression ^23–25, 33^. These changes have been associated with assembling discrete “NF-κB factories” as demonstrated using the *SAMD4A* and *EXT1* TRGs as a model ^23^. These represent specialized nuclear sites involving long-range interactions between promoters and enhancers, leading to the co-regulation of multiple genes ^23^. Another study reported that p65/RelA, a component of NF-κB, assembles into nuclear foci by liquid-liquid phase separation at super-enhancers to activate transcription of individual loci ^40^. Thus, gene activation by NF-κB is an example of possibly different mechanisms that have been reported to link genome structure to transcription programs. In this study, we conducted a genome-wide analysis to identify chromatin modules involved in the TNFα-mediated proinflammatory response, leading us to propose the AC/DC model of gene regulation (**Fig. 8B**). This model distinguishes AC and DC modules, identified via the analysis of dense coverage snATAC-seq with the RWireX software. It enables the identification of stochastic co-accessibility patterns and contiguous domains, which would not be possible with population-averaged data as evident from the highly similar pseudo-bulk profiles acquired for the different time points.

Our findings suggest two co-existing transcription compartment architectures. The AC modules are characterized by long-range co-accessibility interactions between promoters and enhancers involving multiple sites. Based on their bimodal frequency distribution, ACs with a low and a high frequency of occurrence were distinguished (**Fig. 3E**). The two groups may reflect the difference between more dynamic enhancer-promoter contacts that occur in a cell type- or state-dependent manner at lower frequency and stable architectural interactions ^41–43^. Interestingly, we detect patterns of anti-correlated metacell co-accessibility boundaries demarcated by sites of highly correlated accessibility that coincided with frequent ACs. These “blue borders” could relate to the stacking of loops/TAD boundaries ^36, 37^. We identified DCs as regions of locally increased contiguous co-accessibility. These domains could be interpreted as the genomic footprint of loci with confined TF mobility ^18–20^ and/or local TF enrichment by phase separation or other mechanisms ^16, 17, 44^ since they display high TF density and cooperative assembly. DCs can be located within TADs but, like ACs, can also form across TAD boundaries, suggesting a TAD-independent additional layer of genome organization possibly relating to nuclear compartments such as nuclear speckles or the nucleolus ^45^. Consistent with this view, TADs and A/B compartments do not reflect binary states but are probabilistic structures ^12^ where TAD substructures can alternate between the A and B compartment ^46^.

A critical functional distinction of ACs versus DCs is reflected in their transcription bursting parameters (**Fig. 8A**). AC-driven transcription primarily influences burst frequency, likely due to chromatin looping-mediated enhancer-promoter interactions that facilitate frequent transcription initiation events ^43^. Interestingly, AC modules displayed anti-correlated expression of clustered TRGs, arguing for switching of a given enhancer between different promoters, as opposed to complexes where the same enhancer simultaneously drives two genes (**Fig. 6C**). Conversely, co-expression was higher in DC-type TRG clusters. Here, regulation was predominantly mediated by changes in burst size, which can be attributed to locally increased transcription factor occupancy at multiple sites within these contiguous domains. This conclusion aligns well with the mechanism of transcription bursting proposed for a *Gal4* TF cluster ^47^.

In summary, we demonstrate the co-existence and functional impact of two different chromatin module types using TNFα-stimulated HUVECs as a prototypic cellular system. Our AC/DC model reconciles observations from sequencing-based studies and fluorescence microscopy experiments. ACs align with long-range chromatin interactions detected in sequencing data, while DCs correspond to local transcription factor enrichment and align with findings from microscopy-based studies. By identifying integrated AC/DC modules, we demonstrate that these different regulatory mechanisms can act either separately or coexist and cooperate to direct transcriptional responses. The differences in the bursting kinetics could be particularly beneficial for a precise and TRG cluster-specific control of the timing and magnitude of the inflammatory gene expression response. By employing the different chromatin modules separately or in combination, cells could balance speed, precision, and flexibility in transcriptional responses, adapting to diverse physiological demands and environmental cues. The approach and RWireX data analysis framework introduced here extend their potential applications beyond our specific biological system. This is illustrated for other cellular systems and perturbations in human and mouse cell types by the examples given in **Fig. S9**. The application of RWireX to identify DC modules in a mouse model for chronic lymphocytic leukemia (**Fig. S9 C, D**) is particularly noteworthy. In the context of previous findings ^39^, which demonstrate the suppression of malignant B cell proliferation by T-bet (a T-box transcription factor), it illustrates how the AC/DC model could also provide new avenues for exploring dysregulation in disease states. We conclude that the findings from our genome-wide co-accessibility analysis reflect general features of eukaryotic transcriptional regulation. Accordingly, we anticipate that further applications of the framework introduced here will affirm the AC/DC module types in diverse cellular responses while providing insights into the underlying regulatory mechanisms that become activated.

## Acknowledgments

We thank Katharina Bauer, Norbert Mücke, Caroline Knotz, Michele Bortolomeazzi, Lukas Frank and Maiwen Caudron-Herger for help and Oliver Stegle, Carl Herrmann, Daniele Tavernari and the members of the Rippe lab for discussions. The authors are grateful to the DKFZ Genomics and Proteomics and Omics IT and Data Management Core Facilities for sequencing and data management services. This work was supported by German Research Foundation (DFG) grants RI 1283/17-1 within SPP2202 to KR, PA2456/11-2 with SPP2202 and INST186/1479-1 within SFB1565 to AP and DFG FOR2674 (Z01) to JPM. Data storage at SDS@hd was funded by the Ministry of Science, Research and the Arts Baden-Württemberg (MWK) and the DFG through grants INST 35/1314-1 FUGG and INST 35/1503-1 FUGG.

## Methods

### Cell culture and TNFα treatment

HUVECs from pooled donors (Lonza, cat. #00191027, lot #18TL232828) were cultured in endothelial basal medium (EBM-2) with supplements (Lonza, cat. #CC-3162). Low passage number cells were seeded in 18-well μ-slides (Ibidi), starved for 20-24 h in EBM + 0.5% FBS before treatment, and then induced with 10 ng/mL human TNFα (PeproTech, cat. #300-01A) for 0, 30 and 240 min. Cells were then fixed with 4% PFA and stored at 4 °C in PBS (Sigma-Aldrich). For scRNA-seq and snATAC-seq, the time course was conducted in three independent replicates, each starting with different aliquots of HUVEC cells.

### Single-cell sequencing data acquisition

The scRNA-seq libraries were prepared according to the Chromium Next GEM Single Cell 5’ (dual index) protocol v2 from 10x Genomics (Pleasanton, USA). The snRNA-seq libraries were prepared on 384-well plates with the SMART-seq 2.5 protocol as described previously ^48^. The snATAC-seq data were prepared using our improved TurboATAC protocol, which increases Tn5 integration efficiency with the Chromium Next GEM Single Cell ATAC kit v2 from 10x Genomics (Pleasanton, USA) ^29^. Simultaneous 5’ RNA and ATAC libraries from the identical nuclei were prepared according to the Chromium Single Cell Multiome ATAC and Gene Expression protocol v1 from 10x Genomics (Pleasanton, USA). Multiplexed library pools were generated at 2-10 nM concentration of each library. They were paired-end sequenced on a NovaSeq 6000 system (Illumina, San Diego, USA) using S4 flow cells for scRNA-seq and snATAC-seq, S1 flow cells for multiome snRNA-seq and SP flow cells for multiome snATAC-seq libraries. The snRNA-seq libraries were sequenced paired-end on two flow cell lanes of an Illumina NextSeq 550 system (Illumina, San Diego, USA) with 25 and 50 bp read lengths. The UMI sequences were provided as the first eight bases of read 1.

### Preprocessing and basic analysis of scRNA-seq data

Processing of scRNA-seq data was conducted with Cell Ranger (10x Genomics, Pleasanton, USA) including introns and using the provided human GRCh38-2020-A reference. Further processing of data was performed in R with Seurat ^49^. Cells were filtered using a minimal threshold of 100 detected genes, a maximal threshold of 5 percent mitochondrial counts, and a minimal threshold of 5,000 UMI counts. Samples were merged, log normalized, and scaled. Outliers were removed per sample by filtering out cells with more UMI counts than the mean plus twice the standard deviation and outside of plus/minus three times the standard deviation of mitochondrial counts. Single cells were embedded in two-dimensional space using PCA (PC 1-16) and UMAP. Cell cycle stages of single cells were inferred from the expression of cell cycle markers ^50^. Cells in cell cycle stages G2M and S were removed, and G1 cells were embedded in two-dimensional space using PCA (PC 1-20) and UMAP. Differential expression analysis between unstimulated and TNFα stimulated HUVECs was performed for pseudo-bulk counts of samples using DESeq2 ^51^. Differentially regulated genes (TRGs) were identified based on thresholds of absolute log2FCs >1 and adjusted p-values <0.05). The genomic location of TRGs was obtained from Cell Ranger reference arc-GRCh38-2020-A-2.0.0. Genomic distances between TRGs were computed using GenomicRanges and gUtils (see **Supplementary Table S3** for additional references to the software used in our study). TRGs below 500 kb distance were considered a TRG cluster. Further information on scRNA-seq data is provided in (**Supplementary Dataset 1**).

### Preprocessing and basic analysis of snATAC-seq data

The snATAC-seq data were demultiplexed and aligned with Cell Ranger ATAC (10x Genomics, Pleasanton, USA) using the provided human GRCh38-2020-A-2.0.0 reference. Further processing of the data was conducted in R with ArchR ^52^. Cells were filtered using a minimal threshold of 10^4.5^ for the number of unique fragments and a TSS ratio above 7. Cell doublets were removed with Amulet in scDblFinder using a 5^th^ percentile cutoff for significant q-values. Additionally, outliers were removed by filtering out cells with blacklist ratios above the mean plus twice the standard deviation. Single cells were embedded in two-dimensional space using an accessibility matrix of 500 bp tiles, Iterative LSI (LSI components 2-14) and UMAP. Cell cycle stages were inferred by integrating corresponding samples from scRNA-seq data using ATAC gene activity scores and constraining the integration per sample. Cells in G2M and S phase were removed. Single cells were embedded in two-dimensional space using an accessibility matrix of 500 bp tiles, Iterative LSI (LSI components 2-8) and UMAP again. Peak calling of pseudo-bulk accessibility data from all samples was conducted with MACS2 ^53^ in ArchR (extendSummits = 500; reproducibility = 2). Differential accessibility analysis was performed by Wilcoxon test between unstimulated and TNFα stimulated HUVECs (maxCells = 6,000; bias = TSS enrichment, log10(nFrags); normBy = nFrags). Peaks with differential accessibility of an absolute log2FC above 1 and an FDR below 0.05 were considered significant. Further information on snATAC-seq data is provided in (**Supplementary Dataset 1**). For the RWireX plots, pseudo-bulk chromatin accessibility data were normalized by the number of unique fragments.

### Preprocessing and basic analysis of snRNA-seq data

Processing of snRNA-seq data was conducted using the nf-core rnaseq pipeline in Nextflow. Within the pipeline, UMI-tools were used to extract UMI information from read1 and read2 was aligned to the human GRCh38-2020-A reference from Cell Ranger (10x Genomics, Pleasanton, USA) using STAR. Salmon was used to quantify UMI counts in exons at gene level and UMI counts in introns at transcript level. Further processing of data was conducted in R using Seurat ^49^. Cells were filtered using a minimal threshold of 100 detected exon-counted genes, a maximal threshold of 5 percent mitochondrial counts, and filtering out cells with exonic UMI counts above/below the mean plus/minus thrice the standard deviation per sample. Cell cycle S and G2M scores of single cells were inferred from exonic UMI counts of marker genes ^50^, and cells were assigned to G1 if their S and G2M scores were below the sample-specific mean plus standard deviation. Non-G1 cells were removed. Further information on snRNA-seq data is provided in **Supplementary Dataset 1**.

### Multiplex smFISH with the padFISH protocol

The analysis of nascent RNAs was conducted with a multiplex smFISH protocol, termed padFISH, using intronic padlock probes against their cDNA and rolling circle amplification. It combines the hybridization-based in situ sequencing (HybISS) ^54^ and the single-cell resolution in situ hybridization on tissues (SCRINSHOT) methods ^55^. Data were acquired with DNA DAPI staining and detection oligos labeled with Alexa Fluor 488, ATTO 550, Alexa Fluor 647, and Alexa Fluor 750. For the co-expression analysis of the CXCL cluster, all four colors were used. The padFISH data for BIRC2, NFKBIA and SELE bursting kinetics were acquired with three colors (Alexa Fluor 488, ATTO 550 and Alexa Fluor 647). The full padFISH protocol and corresponding image analysis details are described in the **Supplementary Methods**.

### Immunofluorescence

For immunofluorescence (IF), fixed cells were permeabilized in ice-cold 0.2% Triton-X in PBS for 5 minutes, then blocked with 10% goat serum (GS) in PBS for 15 minutes. Incubation with primary antibody mix (Recombinant Anti-NF-kB p65 antibody [E379], ab32536, LOT #GR3275776-15, Abcam) with 10% GS was performed for 1 h. Cells were washed twice with 0.002% NP-40 detergent solution in PBS for 5 minutes. Next, the secondary antibody mix (Goat anti-Rabbit IgG (H+L) labeled with Alexa Fluor 647 (Invitrogen, cat. # A21244, lot #2836809) with 10% GS in PBS was added for 30 minutes. After two 5-minute washing steps in PBS, cells were incubated with 5 μM DAPI in PBS for 15 minutes and then washed three times in PBS. All steps were performed at room temperature. IF samples were stored in PBS at 4°C until imaging.

### Imaging data acquisition

Samples were imaged using an Andor Dragonfly 505 spinning disk confocal unit equipped with a Nikon Ti2-E inverted microscope and a Plan Apo 60x/1.40 oil objective or a 100x CFI SR HP Plan Apochromat Lambda S silicone immersion objective. Multicolor images were acquired for DAPI (λ_ex_ = 405 nm, λ_em_ = 445±23 nm), Alexa Fluor 488 (λ_ex_ = 488 nm, λ_em_ = 521±19 nm), ATTO 550 (λ_ex_ = 561 nm, λ_em_ = 594±21.5 nm), Alexa Fluor 647 (λ_ex_ = 637 nm, λ_em_ = 685±23.5 nm) and Alexa Fluor 750 (λ_ex_ = 730 nm, λ_em_ = 809±45 nm). All images were recorded in Imaris format at 16-bit depth and with 1024x1024 pixel dimensions (pixel size: 0.217 μm or 0.1204 μm) using an iXon Ultra 888 EM-CCD camera. Tiles were recorded as z-stack of 10μm thickness with a step size of 0.4 μm (26 frames) and with 10% overlap.

### Image analysis

For the preprocessing of raw images (Imaris format) and metadata, image stacks were first transformed into maximum projected TIF files. Next, a custom script was used to perform flatfield correction, chromatic aberration correction, and stitching (via Grid/Collection Stitching FIJI plugin). Stitched images were used as input in all subsequent analyses. Nuclei segmentation on DAPI images was performed with Cellpose 2 ^56^ using the pre-trained cyto model with diameters 150 and 200 for 60x and 100x objectives, respectively. Next, cell nuclei at the image’s borders or that displayed overexposure in individual channels were filtered out in R before further analysis. Individual channel images and Cellpose nuclear masks were used as input in R to quantify image features in regions corresponding to nuclear masks using the function *quantNuclei*. For IF and padFISH transcriptional bursting analysis, we computed the sum of fluorescence intensities in each nucleus. Further details are described in the **Supplementary Methods** for the padFISH co-expression analysis at the CXCL cluster. Custom scripts for image analysis are available at https://github.com/RippeLab/padFISH.

### Co-expression analysis of scRNA-seq and padFISH data

Co-expression was computed between all isolated TRGs and TRGs of the same TRG cluster from scRNA-seq data. Spearman correlation coefficients of TRG UMI counts were calculated across single cells per sample. TRGs with expression in less than 10% of cells were removed from the analysis for each sample separately. The mean Spearman correlation coefficient from replicates was used as a co-expression value between two TRGs. The overall co-expression of TRG clusters was computed using mean Spearman correlation coefficients of all TRG combinations within. Co-expression patterns in the CXCL TRG cluster were determined for each cell from scRNA-seq and PadFISH. In scRNA-seq, TRGs CXCL1, CXCL2, CXCL3, and CXCL8 were considered as expressed when showing at least one UMI count in a cell. For padFISH, fluorescence intensity thresholds were defined from the minimum of the bimodal intensity distribution and adjusted by visual inspection of each channel in a pre-segmented region obtained from the co-localization of the 4 TRGs (**Supplementary Methods**). Fractions of cells or subcellular loci with different CXCL co-expression patterns were quantified for each sample from scRNA-seq and PadFISH.

### Co-accessibility analysis with RWireX

Single-cell and metacell co-accessibility were computed for replicates of snATAC-seq data using RWireX. Details of the co-accessibility analysis with RWireX are described in the **Supplementary Methods**. The resulting AC and DC features at TRGs were annotated using GenomicRanges. AC start and end peaks and DCs at TRG promoters, defined as ±500 bp around the TSS, were quantified. The number of DCs was binarized, while the number of ACs was log10 transformed with a pseudo-count of 1. TRGs were clustered by min-max normalized AC and DC features (5 clusters from ward.D clustering) and visualized by heatmap. Clusters were termed by the prevalent feature and used to annotate TRGs as AC-, DC- or AC/DC-driven or not assigned (NA). Next, the AC and DC scores of each TRG cluster were computed from the AC/DC annotation of the TRGs in the cluster as *S_AC_* = (Σ*TRG_AC_* + Σ*TRG_AC/DC_*)/*n* and *S_DC_* = (Σ*TRG_DC_* + Σ*TRG_AC/DC_*)/*n* with *n* being the total number of TRGs in the cluster. Genomic TRG clusters were then assigned as (i) AC for *S*_AC_ ≥ 0.5 and *S*_DC_ < 0.5; (ii) DC for *S*_AC_ < 0.5 and *S*_DC_ ≥ 0.5, (iii) AC/DC for *S*_AC_ ≥ 0.5 and *S*_DC_ ≥ 0.5 and (iv) NA for *S*_AC_ < 0.5 and *S*_DC_ < 0.5.

### Analysis of TF binding

The TF binding activity score *TF*_score_ was calculated from pseudo-bulk footprints of snATAC-seq replicates using Tobias in Python ^34^ with Homer universal motifs from chromVARmotifs in 1 kb ATAC peaks. Increased TF binding between unstimulated and TNFα stimulated HUVECs was determined by log2FC of the number of TF-bound sites (replicate average) with a log2FC > 0.1 threshold. Genome-wide and region-specific (local DC or non-DC background) TF footprints were visualized per sample using pseudo-bulks of 1,000 cells without normalization and a smoothing window of 20 in ArchR. To assess differential TF binding between genomic regions, we compared the *TF*_scores_ in DCs and non-DC regions (global background) for TNFα responsive TFs. Differential genomic TF binding was calculated as *log*2*FCTF_score_* = *log*2(〈*TF_score,DC_*〉) − *log*2(〈*TF_score,back_*〉) and using a one-sided Wilcoxon test per DC comparing the *TF*_scores_ in the respective DC and the global non-DC background *TF*_scores_. Results from replicates were combined by meta-analysis with Fisher’s method using poolr and averaging of log2FCs. TFs with differential genomic binding p-value below 0.05 and log2FC above 1 were considered significantly locally enriched in the respective DC. DCs with significant local enrichment of TF binding activity were visualized by heatmap, clustering DCs by summed TF family enrichment with ward.D2.

### Preprocessing and data analysis of multiome snRNA- and snATAC-seq

Multiome snRNA- and snATAC-seq data were processed with Cell Ranger ARC (10x Genomics, Pleasanton, USA), including introns, using the provided human GRCh38-2020-A reference. Further processing of data was conducted in R using Seurat and ArchR. For RNA data, high-quality cells were selected using a minimal threshold of 5,000 UMI counts and minimal and maximal thresholds of 5 and 40 percent mitochondrial counts. Outliers were removed per sample by filtering out cells with UMI counts above the mean plus twice the standard deviation. Samples were merged, log normalized, and scaled. Cell cycle stages of single cells were inferred from the expression of cell cycle markers ^50^, and cells in G2M and S were removed. For ATAC data, high-quality cells were selected using minimal thresholds of 10^3.5^ unique fragments and a TSS enrichment score of 7. Cell doublets were removed using Amulet in scDblFinder. Additionally, outliers were removed by filtering out cells with unique fragments above 30,000 and blacklist ratios above the mean plus twice the standard deviation. A mixed cluster composed of 86 cells from all conditions was excluded. Finally, high-quality cells from both ATAC and RNA were selected. Further information on multiome snRNA- and snATAC-seq data is provided in (**Supplementary Dataset 1**).

Gene expression in high-quality cells was quantified using intronic and exonic UMIs in Ensembl annotated genes for the Cell Ranger ARC human GRCh38-2020-A reference. TRG expression was correlated to ATAC features (ACs and DCs) using Spearman correlation from SciPy in Python. For ACs, chromatin accessibility in high-quality cells was quantified using insertions in peaks from snATAC-seq data. Accessibility counts of each link’s start and end peaks were multiplied to obtain AC activities per cell. TRG expression was correlated to the activity of ACs at the TRG’s promoters. For DCs, chromatin accessibility in high-quality cells was quantified using insertions within the whole domains. TRG expression was correlated to the accessibility of DCs comprising the TRG promoter. For exemplary regions, TRG expression and chromatin accessibility were visualized using a heat map. Accessibility was quantified within 2 kb bins of the region. Cells were hierarchically clustered by TRG expression using SciPy with Euclidean distances and average linkage.

### Transcriptional burst kinetics from snRNA-seq and padFISH data

A two-state model of transcription was applied that yielded the burst frequency rate *k*_on_ and the burst size from the ratio of *k*_syn_/*k*_off_ according to the mechanism depicted in **Fig. 8A** ^57^. To compute these parameters, we used intronic UMI counts of TRG transcripts from snRNA-seq data at single-cell resolution according to the equation. Only TRG transcripts with intronic UMI counts in at least 5% of cells across all treatment conditions and showing the same direction of TNFα regulation as gene-level TRGs in the scRNA-seq replicate data were used. Capture efficiency was estimated from total transcriptome UMIs per sample, assuming 20% of 500,000 mRNA molecules/cell in the nucleus (0.33 for 0 min; 0.36 for 30 min; 0.21 for 240 min). Weighted averages of transcript-level burst sizes and frequencies were calculated per treatment condition to obtain TRG-level burst kinetics.

Transcriptional burst kinetics of NFKBIA, SELE, and BIRC2 were inferred from padFISH data in two replicates following the same model. Thresholds for active transcription were determined from the minima of bimodal nuclear fluorescent intensity distributions per TRG and replicate. Cells with nuclear fluorescent intensities below these thresholds were considered not actively transcribing the respective TRG. A transcript detection efficiency of 0.35 was estimated for padFISH ^58^. Average transcript lengths were used per TRG to approximate transcription time. For comparison, burst sizes and frequencies were scaled from zero to one for snRNA-seq and padFISH, respectively.

### Analysis of 3’ RNA, H3K27ac ChIP and Hi-C bulk sequencing data

The differential expression analysis between unstimulated and TNFα stimulated HUVECs by bulk RNA-seq was conducted, reanalyzing the data from ref. ^26, 27^ on the hg38 reference genome using HISAT2 ^59^. Gene counting was performed using HTSeq ^60^, and TMM normalization was carried out to adjust for differences in library sizes across samples. Differential expression was analyzed with NOISeq ^61^, using five technical replicates for each condition. Genes with a differential expression probability ≥ 0.8 were considered significant. H3K27ac ChIP-seq with two H3K27ac antibodies (Active Motif, 39133; Diagenode, C15210016) was conducted as described previously ^27^. Sequencing reads were aligned to the hg38 reference genome using Bowtie2 ^62^ and peak calling was conducted as described previously ^63^. Peaks with a FDR ≤0.01 and a peak height ≥20 were selected. GenomicRanges was used to compute the overlap of H3K27ac peaks with TRGs, ATAC peaks, and co-accessibility features. The contact matrices from the Hi-C-seq data of unstimulated HUVECs were from ref. ^28^. They were converted from hic to cool format using hic2cool and lifted to the hg38 genome with HiCLift. The final contact matrices were used as input for Arrowhead ^28^ to call TADs at 25 kb resolution. Overlap of TADs with TRG clusters and co-accessibility features was assessed using GenomicRanges. Balanced counts were extracted from cool files using cooler with parameters –join -b –balanced ^64^. Contact counts between AC-linked genomic sites were retrieved by mapping ATAC peaks to 10 kb bins of the Hi-C contact matrix using GenomicRanges. Contact matrices of exemplary regions at 10 kb resolution were visualized as heatmaps using plotgardener in R. Pile-up plots of chromatin contacts at TADs and ACs were created with coolpup.py ^65^.

## Data availability

The datasets that can be directly downloaded with the manuscript are listed in **Supplementary Table S2**. An overview of the data from this manuscript at public repositories is given in **Supplementary Table S4**. Sequencing and imaging data are available from the following locations: The single-cell sequencing data and bulk H3K27ac and RNA-seq are available at GEO accession number GSE273430 with separate accession numbers for the individual datasets of scRNA-seq (GSE273426), snATAC-seq (GSE273428), snMultiome-seq (GSE273429) and snRNA-seq (GSE273427). The Hi-C-seq from ref. ^28^ are archived at GSE63525. The padFISH source images have been deposited in the BioImage Archive under accession number S-BIAD1294 (https://www.doi.org/10.6019/S-BIAD1294). Custom analysis software tools are available on Github at https://github.com/RippeLab/RWireX with test data deposited at https://www.doi.org/10.5281/zenodo.13142236 (RWireX) and https://github.com/RippeLab/padFISH (padFISH). Other data analysis software used in our study is listed in **Supplementary Table S3**.

