## Supplementary Material for "Two distinct chromatin modules regulate proinflammatory gene expression"

#### Inventory of Supplementary Information

##### Supplementary Methods

padFISH multiplex smFISH of nascent RNAs
Co-accessibility analysis with RWireX

##### Supplementary Figures

Figure S1. TNFα-induced gene expression and chromatin accessibility.
Figure S2. Genomic location and co-expression of TRGs.
Figure S3. Single cell co-accessibility analysis with RWireX.
Figure S4. Metacell co-accessibility analysis with RWireX.
Figure S5. Local TF binding activity.
Figure S6. Features of AC, DC and AC/DC chromatin modules.
Figure S7. Hi-C chromatin contacts at AC and DC chromatin modules.
Figure S8. Bursting kinetics of TRGs in AC, DC and AC/DC chromatin modules.
Figure S9. DCs and ACs in mouse cells under perturbation.

##### Supplementary Tables

Table S1. padFISH of CXCL intronic transcripts.
Table S2. Inventory of supplementary data sets associated with this manuscript.
Table S3. Data analysis packages and software from external sources.
Table S4. Inventory of data sets deposited at external repositories.

##### Supplementary References

### Supplementary Methods

#### padFISH multiplex smFISH of nascent RNAs

The padFISH method integrates two protocols: hybridization-based in situ sequencing (HybISS) [1] and single-cell resolution in situ hybridization on tissues (SCRINSHOT) [2]. In our study, we employed this technique to trace nascent RNAs using intronic padlock probes (PLPs) targeting their cDNA and to detect their rolling circle amplification (RCA) product.

Experimental padFISH protocol
Cells fixed and stored in PBS underwent permeabilization with 0.1 M HCl for 3 minutes at room temperature (RT), followed by three 1-minute washes in PBS/Tween 0.05%. Subsequently, the cells were exposed to a blocking mix for 30 minutes at RT and washed twice for 1 minute with PBS/Tween 0.05%. For PLP hybridization, we applied a solution containing 20% formamide and 50 nM of each probe, then incubated the slides for 15 minutes at 55 °C followed by 2 hours at 45 °C. After hybridization, cells underwent three 10-minute washes with 10% formamide in 2x SSC at 45 °C, and then three 1-minute washes with PBS/Tween 0.05%. To ligate PLPs, the sample was incubated with 0.5 U/ml of SplintR ligase (New England Biolabs) for 15 minutes at 25 °C. After ligation, cells were washed twice for 1 minute in PBS/Tween 0.05%. We then performed RCA by incubating the sample overnight at 30 °C with NxGen Phi29 DNA polymerase (LGC Biosearch Technologies) and an RCA primer (100 nmol DNA oligonucleotide from Integrated DNA Technologies (IDT)). After RCA, cells were washed three times for 1 minute in PBS/Tween 0.05% and subsequently fixed in 4% PFA for 15 minutes at RT. Following another set of three washing steps in PBS/Tween, cells were incubated with 65% formamide in 2x SSC at 30 °C, then underwent the usual PBS/Tween washes and an additional washing step in 2x SSC. Bridge probe (BP) and detection oligonucleotide (DO) hybridization were performed according to the L-probes HybISS Protocol [1]. For BP hybridization, we added a solution of 4x SSC, 40% formamide, and each barcode at a final concentration of 0.1 mM to the wells and incubated them for 1 hour at RT. Excess probes were removed by washing with 2x SSC. DO hybridization using 0.1 mM of each barcoded dye was carried out simultaneously with DAPI staining for 1 hour at RT. Excess DOs were removed by washing three times with PBS and cells were stored in PBS at 4 °C until imaging.

padFISH probes
Based on the scRNA-seq analysis, we designed PLPs for padFISH targeting selected TNFα regulated genes (TRGs) using a combinatorial approach that combined the SCRINSHOT probe design criteria [2] with the bar/detection scheme of the HybISS method [3]. The PLPs were designed to bind intronic regions of CXCL1, CXCL2, CXCL3, CXCL8, BIRC2, NFKBIA and SELE for nascent RNA detection. We determined the number of PLPs per gene through previous tests across replicates, using 3 (CXCL3), 4 (CXCL1, CXCL2, CXCL8), 5 (BIRC2) and 7 (NFKBIA, SELE) probes. All PLPs, BPs and DOs were obtained from IDT. PLPs were ordered as DNA oligonucleotides with 5'-phosphate modification and 20 nmol synthesis scale (200 µM in IDTE buffer pH 8.0). BPs were synthesized as 25 nmol DNA oligonucleotides (200 µM in IDTE buffer pH 8.0). DOs were acquired as 1 µmol DNA oligonucleotides with 5'-modifications of Alexa Fluor 488, ATTO 550, Alexa Fluor 647, and Alexa Fluor 750.

Co-expression analysis of nascent RNAs
For the padFISH co-expression analysis of the CXCL cluster, individual channel images (488 nm, 651 nm, 637 nm, 730 nm) were used to detect the co-expression locus across the 4 TRGs of

interest. Z projection (SUM function) was applied to all channels to identify regions with co-localization of at least two channels. The Z-projected image was filtered with Gaussian Blur 2.0 and then used to create a segmentation mask of the co-expression locus in ilastik (segmentation workflow: pixel classification). In FIJI, the output image was binarized and filtered (Gaussian blur 1.5). An intensity threshold (Otsu, 230/255) was established, and co-expression masks were saved as regions of interest (ROIs) using the particle analyzer tool (size: 0-Infinity; circularity: 0.00-1.00). FIJI's Multimeasure function was applied to the pre-selected ROIs to quantify area, mean gray value, and center of mass across all channels (including the DAPI channel as input).

Next, combined ROIs were converted into masks. We then used nuclear masks, co-expression masks, and individual channel images corresponding to the target TRGs as input in R to compute co-expression. The co-expression masks were assigned to nuclei using our custom function *quantNuclei* [4]. For the analysis, only nuclear masks containing a minimum of one and a maximum of two co-expression masks were selected. Quantification was performed exclusively across the co-expression locus masks. We further filtered out cells from the analysis if the subcellular co-expression masks' area and sum of intensities represented outliers from the overall population. Minimum fluorescence intensity thresholds for CXCL1, 2, 3, and 8 were defined from the minimum bimodal intensity distribution and adjusted based on visual inspection of each channel in the co-expression mask. Finally, the expression pattern of target genes was binarized (1 for active, 0 for inactive).

#### Co-accessibility analysis with RWireX

The co-accessibility analysis was conducted with the RWireX software, which is based on our previous work [5, 6] and is available at https://github.com/RippeLab/RWireX. It identifies chromatin regions simultaneously open and accessible in the same cells or metacells from snATAC-seq data. RWireX is implemented as an extension of the ArchR package [7]. It computes Pearson correlation coefficients across different cell populations and at varying levels of resolution using two distinct workflows, which are illustrated in the scheme below. We employed this approach to identify both autonomous links of co-accessibility (ACs) and domains of contiguous co-accessibility (DCs).

The "single cell co-accessibility" workflow identifies ACs from stochastic accessibility changes in 1 kb ATAC peaks. It employs a homogeneous population of single cells as input, e. g., in our case, the separate 0 min, 30 min, and 240 min time points of TNFα treatment. Pearson correlation coefficients between two peaks are evaluated against a local background model. Background co-accessibility is calculated from the 99th percentile derived from accessibility matrices per chromosome that are shuffled over cells and peaks. The stability of ACs is determined from their prevalence in the single-cell population by computing the average percent of accessible cells for the linked peaks.

The "metacell co-accessibility" workflow identifies DCs from accessibility changes in 10 kb genomic tiles and utilizes profiles from aggregated cells (metacells) with similar chromatin accessibility profiles as input. This workflow requires input from a heterogeneous cell population. Therefore, different cell types or states are jointly used, e.g., in our case, the combined 0 min, 30 min, and 240 min time points of TNFα treatment. This approach enables the identification of broader genomic patterns of depleted or enriched co-accessibility along the genomic coordinate, driven by heterogeneous cell states at the TNFα treatment time points. The efficacy of the RWireX analysis improves with higher snATAC-seq coverage. This parameter was particularly high in the dataset studied here with ~150,000 unique fragments/cell. Based on our experience, snATAC-seq data with 20,000-30,000 unique fragments/cell are also suitable for analysis with RWireX.

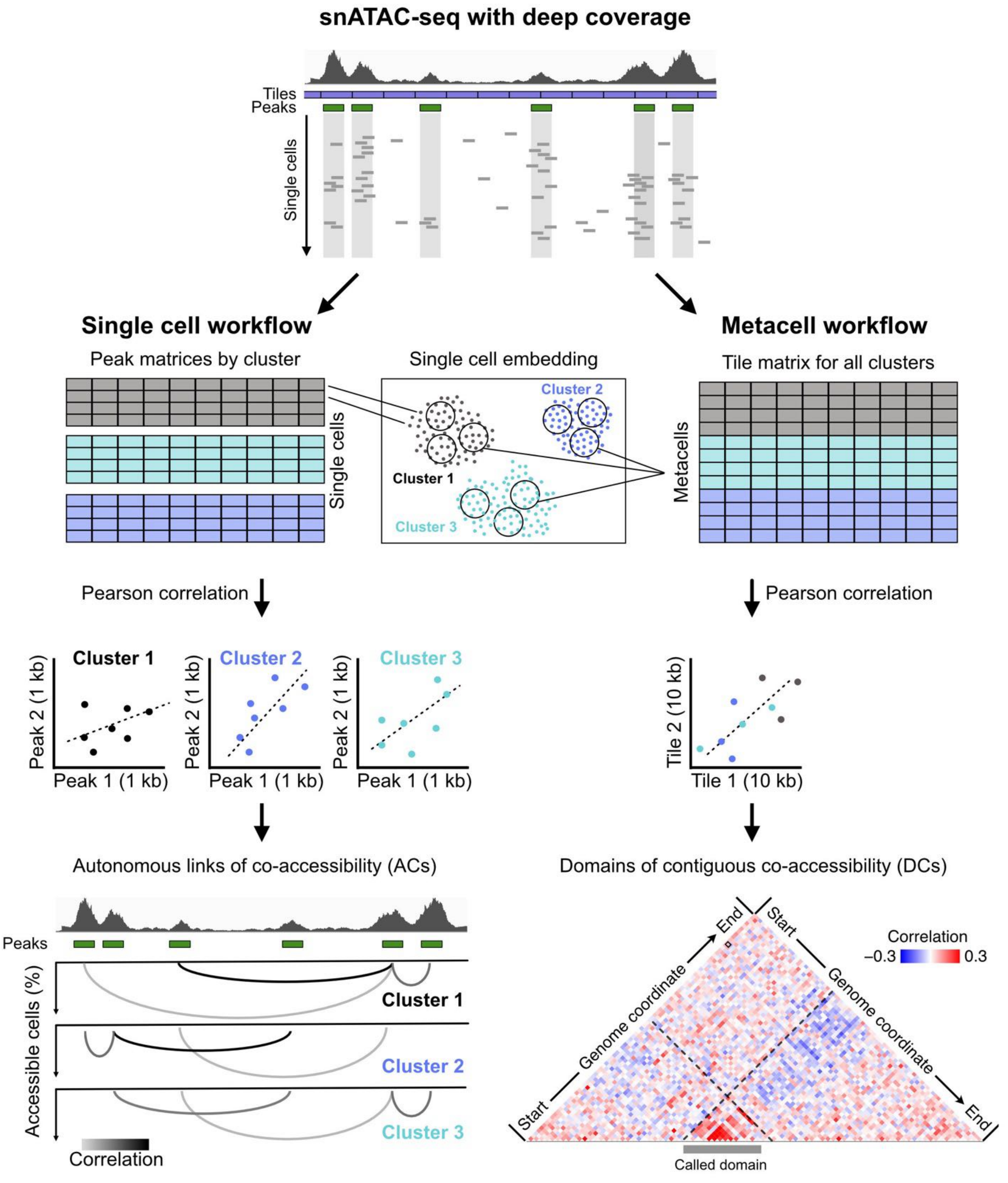


<u>Single cell co-accessibility</u>

The co-accessibility analysis with RWireX conducted here utilized 1,000 cells per sample, all in the G1 phase of the cell cycle and with similar numbers of unique fragments per cell. Single cell co-accessibility was computed from a continuous accessibility matrix of ATAC peaks within 1 Mb. Co-accessible links were filtered using sample-specific background co-accessibility cutoffs (0.083-0.088), a minimal cutoff of 5% accessible cells, and by retaining only positively correlated links. This process resulted in a set of ACs for each sample. Consensus ACs for each treatment condition were obtained by selecting links detected in at least two replicates, and averaging Pearson correlation coefficients and percent accessible cells from these replicates. The reproducibility of ACs was assessed between replicates by computing the percent of consistent links among all links of the samples. TRGs, H3K27ac peaks, and TADs were annotated at the start and end peaks of ACs using GenomicRanges [8].

Metacell co-accessibility

Metacells were formed from unique sets of 10 cells each, with 10% of cells excluded to prevent forced aggregation of dissimilar cells. For each replicate, metacell co-accessibility was computed across treatment conditions using a continuous accessibility matrix of 10 kb genomic tiles within 2 Mb regions. Consensus metacell co-accessibility was obtained by averaging the Pearson correlation coefficients of replicates and visualized as heatmaps of co-accessibility matrices using plotgardener [9]. DCs were determined from positive co-accessibility matrices (consensus and individual replicates) using SpectralTAD [10]. The algorithm was run with three levels of domain hierarchy for small domains (minimal domain size of 20 kb, window size of 200 kb) and large domains (minimal domain size of 200 kb, window size of 2 Mb) separately. The average of Pearson correlation coefficients within determined the overall co-accessibility of domains. Lower cutoffs from 90th co-accessibility percentiles were applied to small and large domains to filter for domains of locally enriched co-accessibility. The reproducibility of DCs between replicates was assessed by computing the percent of bp overlap between domains using GenomicRanges. Differential accessibility analysis of domains was conducted by Wilcoxon test between unstimulated and TNFα stimulated HUVECs (maxCells of 6,000; bias correction by TSS enrichment, log10(nFrags); normalization by nFrags). Domains with differential accessibility of FDR below 0.05 were considered significant. TRGs, ATAC peaks, H3K27ac, and TADs were annotated in DCs using GenomicRanges.

Metacell co-accessibility of mouse data

For the analysis of co-accessibility in mouse cells under perturbation, we utilized snATAC-seq data from mouse embryonic stem cells (ESCs) and embryonic fibroblasts (MEFs) treated with IFNβ [6], as well as from TCL1 mouse models upon Tbx21 knock-out [11]. Comparable cell numbers and numbers of unique fragments per cell were used for ESCs and MEFs (2,700 cells) and wild-type and Tbx21-knockout TCL1 cells (1,000 cells). Metacell co-accessibility was computed using continuous accessibility matrices of 10 kb genomic tiles within 2 Mb regions. Metacells were generated from 10 unique cells each, omitting 10% of cells to prevent forced aggregation. Co-accessibility matrices were visualized as heatmaps using plotgardener. For ESCs and MEFs, we annotated interferon-stimulated genes (ISGs) and STAT1/2 bound sites from bulk RNA- and ChIP-seq data [6]. For TCL1 cells, we annotated T-bet-dependent genes from bulk RNA-seq data [11] and Tbx21 and NF-κB binding motifs from Homer [12].

#### Supplementary Figures

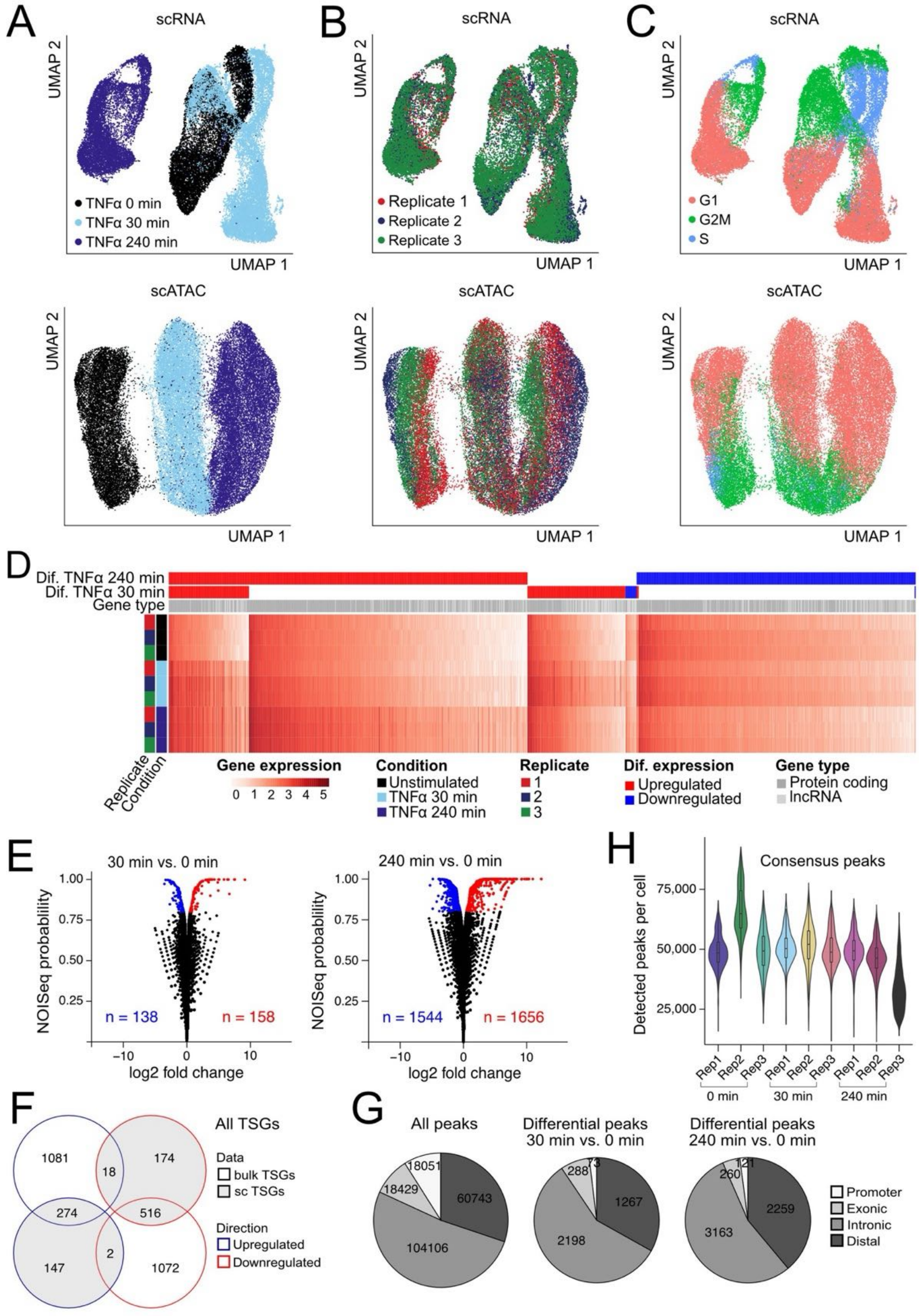


**Supplementary Figure 1. TNFα-induced gene expression and chromatin accessibility**. (**A**) UMAP embeddings of scRNA- (top) and snATAC-seq data (bottom) from three biological replicates with coloring according to TNFα treatment. (**B**) Same as panel A with coloring of biological replicates. (**C**) Same as panel A with coloring of cell cycle states. Only cells in G1 cell cycle state were used for subsequent analyses. (**D**) Expression of TNFα regulated genes (TRGs) with log10 transformed and scaled UMI counts. TRGs are annotated by gene type and differential regulation after 30 and 240 min of TNFα treatment. (**E**) Differential expression after TNFα treatment in 3' bulk RNA-seq data. Significant differential expression is defined as log2 fold change (log2FC) ≥ 1 (up-regulated, red) or log2FC ≤ -1 (down-regulated, blue) and differential expression probability ≥ 0.8. (**F**) Overlap between TRGs from pseudo-bulks of 5' scRNA-seq replicates and 3' bulk RNA-seq. Only protein-coding TRGs are shown. The circle colors indicate the direction of regulation. For TRGs at both time points, 30 min results are considered. (**G**) Number and location of peaks from pseudo-bulk analysis of snATAC-seq. Peaks were classified as promoter, exonic, intronic, and intergenic. Left: All 201,329 ATAC peaks. Right: Differential ATAC peaks after 30 min and 240 min of TNFα treatment.

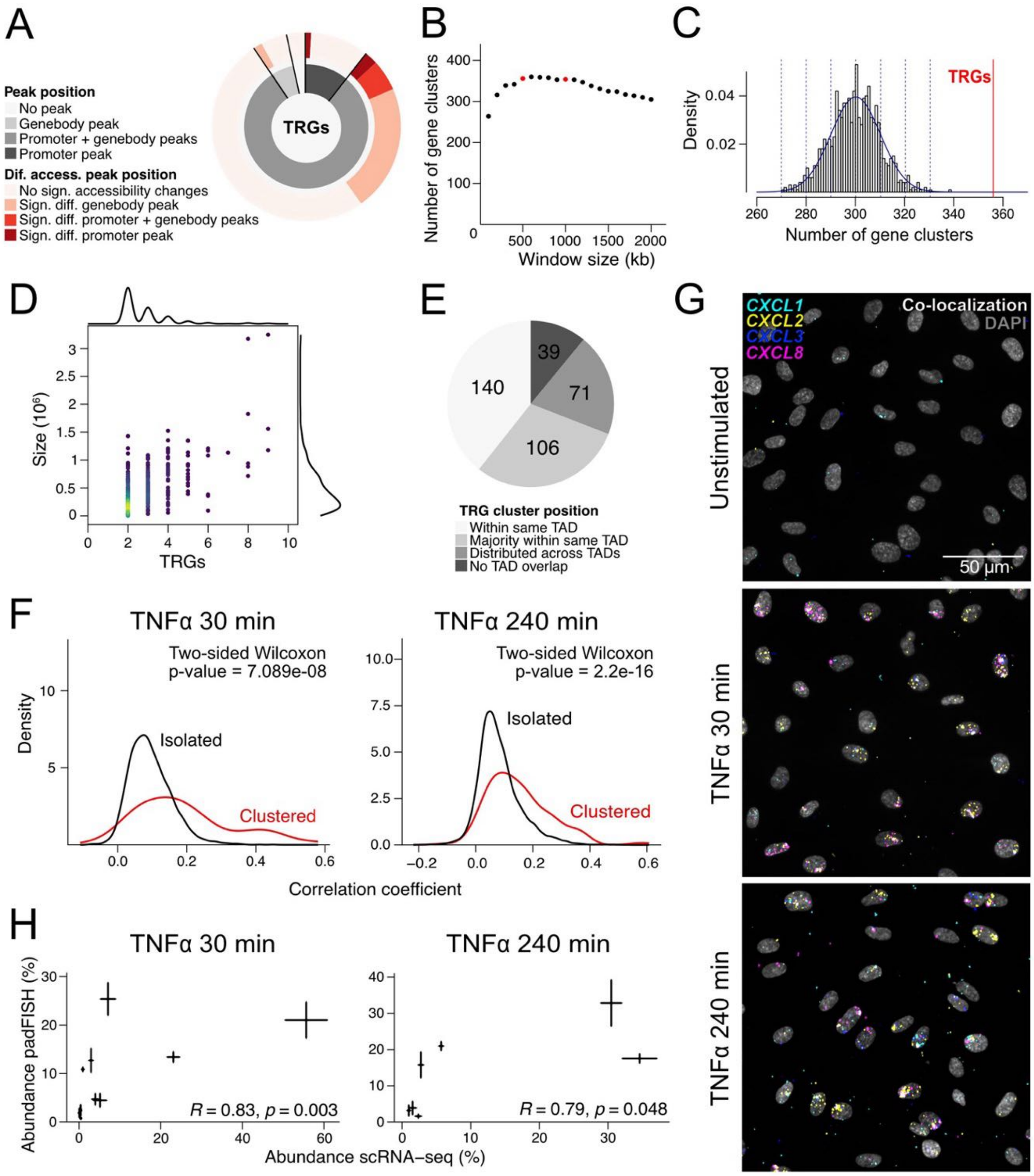


**Supplementary Figure 2. Genomic location and co-expression of TRGs.** (**A**) Proximal regulation of TRG expression by promoter and/or gene body ATAC peaks. The inner wheel shows the fraction of TRGs with ATAC peaks at their promoter, gene body, promoter, and gene body, or none. The outer wheel shows the fraction of TRGs with significantly differential ATAC peaks in the previous groups. (**B**) Number of TRG clusters detected in dependence of distance cutoffs to define TRG neighbors. (**C**) Number of gene clusters from 1,499 genes randomly selected 1,000 times with a distance cutoff of 500 kb for local neighbors. This yielded about 300 clusters on average. The number of 356 TRG clusters detected at this distance is significantly higher and marked by a red line. (**D**) TRG cluster size over the number of TRGs per cluster. The colors of points reflect the density of TRG clusters. Density curves of TRG cluster size and number of TRGs are shown. (**E**) Genomic location of TRG clusters in relation to TADs from Hi-C data of unstimulated HUVECs. TRG clusters were classified as all TRGs within the same TAD, majority of TRGs within the same TAD, TRGs distributed across TADs, and no TAD overlap of TRGs. (**F**) Co-expression of clustered (red) and isolated (black) TRGs after 30 min (left) and 240 min (right) of TNFα-treatment. Average replicate co-expression for upregulated TRGs is shown with p-values from a two-sided, unpaired Wilcoxon test. Clustered TRGs show significantly higher co-expression than isolated TRGs. (**G**) padFISH images of intronic CXCL1 (cyan), CXCL2 (yellow), CXCL3 (blue), and CXCL8 (magenta) expression after 0, 30 and 240 min of TNFα treatment. (**H**) CXCL co-expression patterns from scRNA-seq and padFISH. Error bars display standard errors from triplicates. Spearman correlation coefficients are indicated. Co-expression results from scRNA-seq and padFISH are highly correlated.

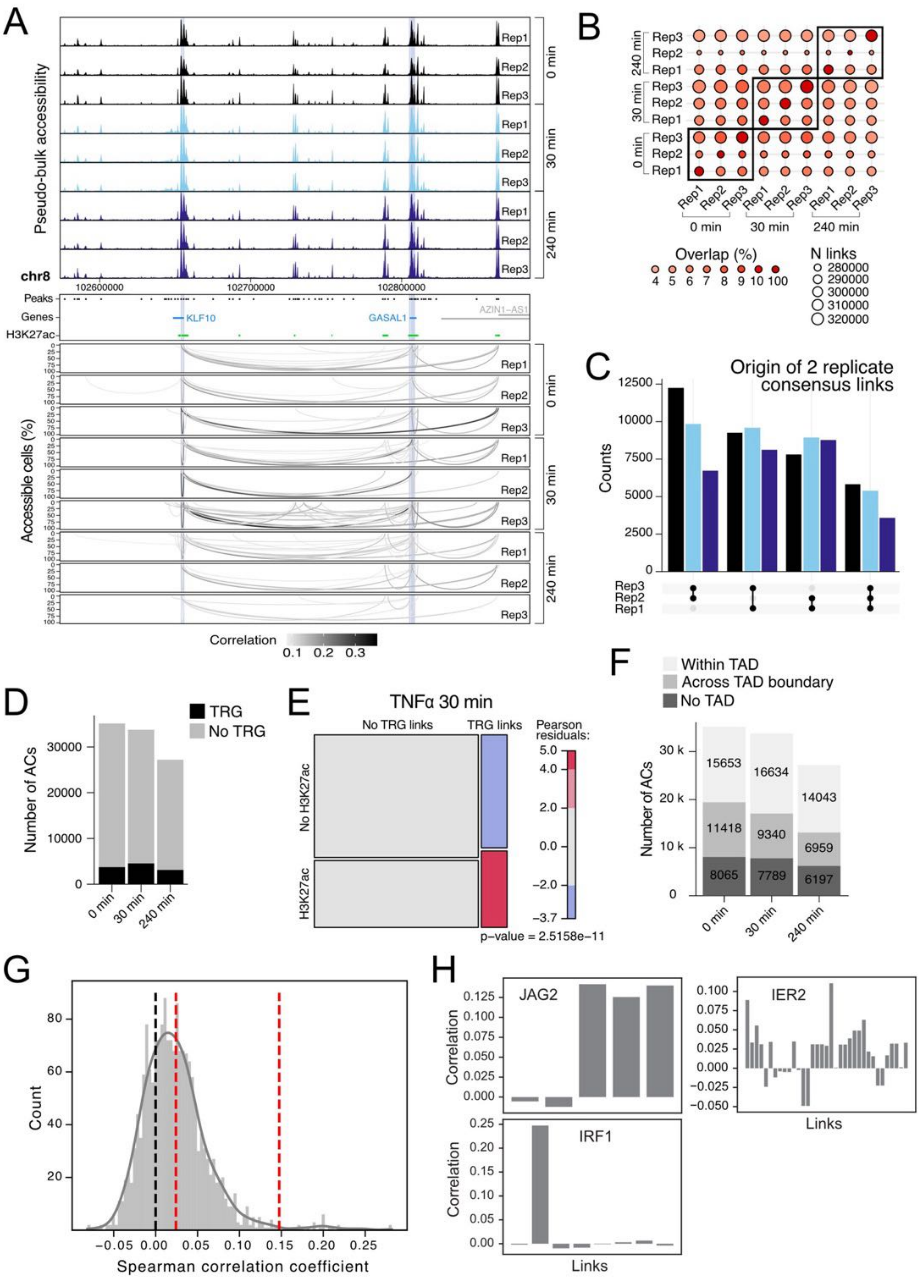

**Supplementary Figure 3. Single-cell co-accessibility analysis with RWireX.** (**A**) Chromatin accessibility and co-accessible links in replicates at the KLF10 and GASAL1 TRG cluster during TNFα induction. Top: aggregated pseudo-bulk chromatin accessibility in three biological replicates. Middle: 1 kb extended pseudo-bulk ATAC peaks (black), genes (gray), TRGs (blue) and 1 kb regions around the TSSs of TRGs (light blue). Bottom: co-accessible links at TRG promoters from single cell co-accessibility of biological replicates. The grayscale and height of loops reflect accessibility correlation and percent accessible cells of linked peaks. (**B**) Reproducibility of co-accessible links between replicates. The size and color of the dots show the total number of links detected in the reference sample and the percent overlap between the samples. Genome-wide co-accessible links show high heterogeneity between biological replicates. (**C**) Number of reproducible co-accessible links in replicates. Reproducible co-accessible links in at least two replicates were used as the consensus set of autonomous links of co-accessibility (ACs). (**D**) Number of consensus ACs at treatment time points at TRGs (black) and non-TRG regions. (**E**) Mosaic plot of ACs with/without TRGs and H3K27ac at their start or end peak. P-value from Chi-squared test is provided. (**F**) Genomic location of ACs in relation to TADs from Hi-C data of unstimulated HUVECs. ACs were classified as within one TAD, across TAD boundary, and no TAD overlap. (**G**) Spearman correlation coefficients of TRG expression and the activity of ACs at the TRG's promoters. Red lines indicate mean as well as mean plus thrice the standard deviation. (**H**) Spearman correlation coefficients of TRG JAG2, IER2 and IRF1 expression and the activity of ACs at the TRG's promoters.

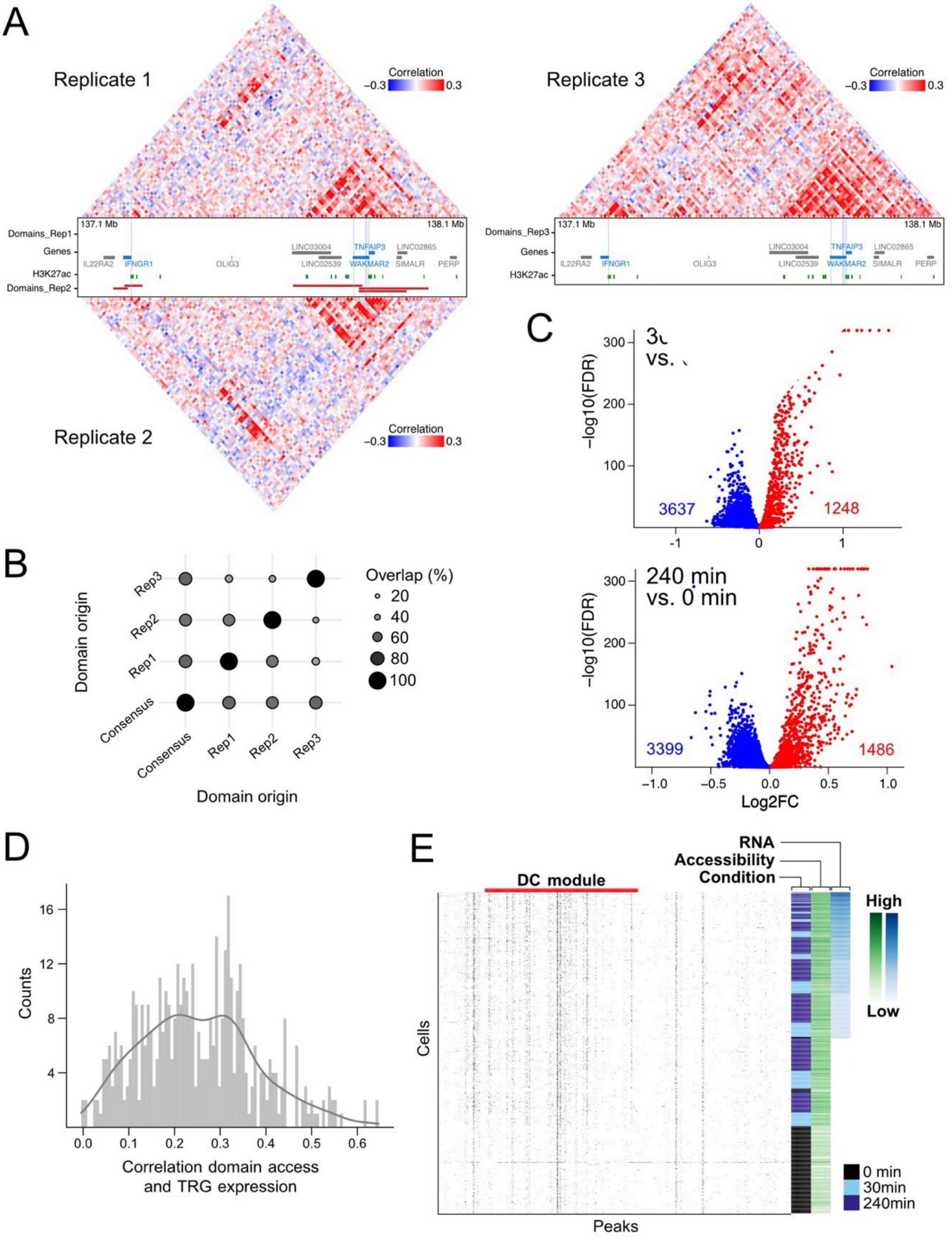


**Supplementary Figure 4. Metacell co-accessibility analysis with RWireX**. (**A**) Chromatin co-accessibility maps at the TNFAIP3, IFNGR1 and WAKMAR2 TRG cluster. DCs from metacell co-accessibility of replicates (black), genes (gray), TRGs (blue) and 1 kb regions around the TSSs of TRGs (light blue) are indicated. (**B**) Reproducibility of co-accessible domains between replicates and consensus DCs. Consensus DCs are computed from the average metacell co-accessibility of replicates. The size and color of the dots reflect the percent of base pair overlap between the domains. (**C**) Differential accessibility in DCs after TNFα treatment of HUVECs across three biological replicates. Differential accessibility is visualized by log2FC and FDR. DCs with FDR below 0.05 are considered significant and marked in red (upregulated) and blue (downregulated). (**D**) Spearman correlation coefficients of TRG expression and the accessibility of DCs comprising the TRG promoters. A positive correlation between DC accessibility and gene expression is observed. (**E**) Expression of TNFAIP3 and chromatin accessibility (2 kb bins) at the TRG cluster of TNFAIP3, IFNGR1 and WAKMAR2. The merged TNFAIP3 DC is depicted in red. Cells are annotated by TNFα treatment condition. The heatmap shows an increased accessibility in the whole DC with high TNFAIP3 expression and a Spearman correlation coefficient of 0.45.

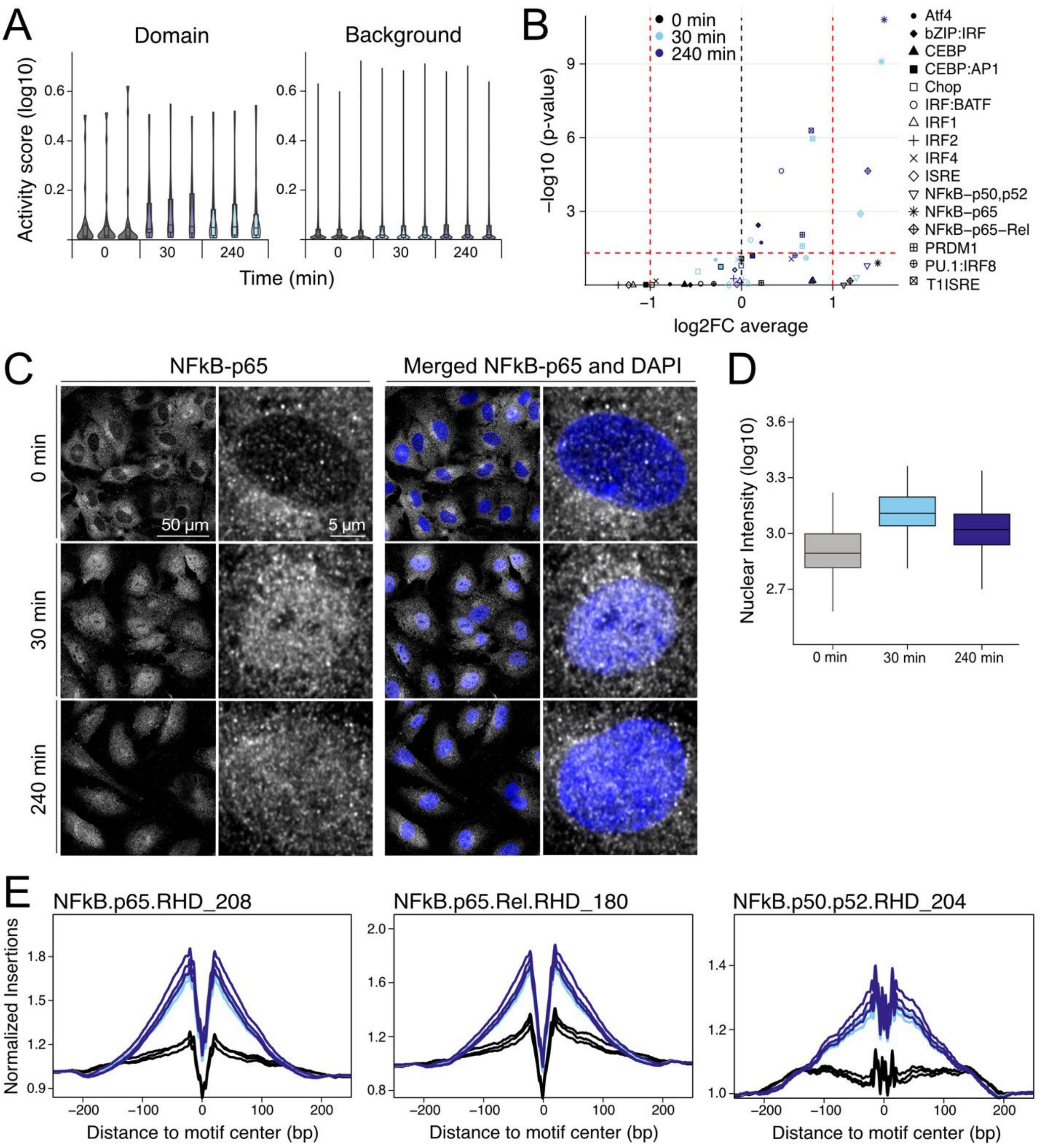


**Supplementary Figure 5. Local TF binding activity.** (**A**) Binding activity scores (log10) of accessible NF-κB/p65 motifs in the merged TNFAIP3 DC. (**B**) Same as panel A with binding activity scores of accessible NF-κB/p65 motifs in genome-wide non-DC regions. (**C**) Differential binding activity in the merged TNFAIP3 DC vs. genome-wide non-DC regions. Differential binding activities across replicates of unstimulated and TNFα-stimulated HUVECs are visualized by average log2FC and p-values from meta-analysis. TFs with absolute log2FC above 1 and FDR below 0.05 are considered significant. Binding activity at accessible NF-κB/p65 and NF-κB/p65/Rel motifs was significantly enriched upon TNFα treatment in the merged TNFAIP3 DC. (**D**) Images of NF-κB IF. Zoom-ins of exemplary cells are shown. (**E**) Nuclear NF-κB signal at different time points quantified from IF images. (**F**) Genome-wide footprints at accessible NF-κB/p65 (left), NF-κB/p65/Rel (middle) and NF-κB/p50/p52 (right) motifs. Three replicates are shown with the different time points indicated by color. NF-κB footprints are present already at the uninduced time point and become stronger upon TNFα treatment.

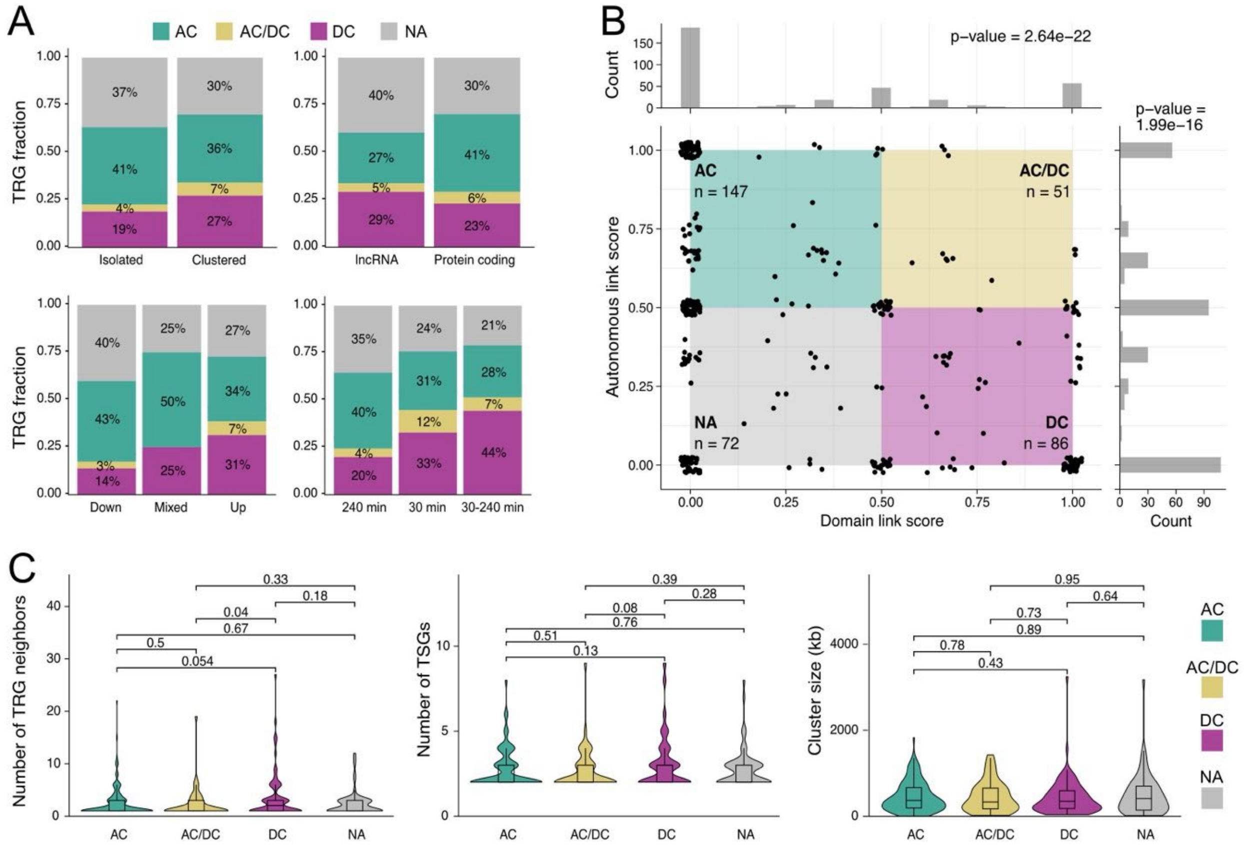


**Supplementary Figure 6. Features of AC, DC, and AC/DC chromatin modules**. (**A**) Proportion of TRGs regulated by AC, DC and AC/DC chromatin modules in clustered and isolated TRGs; lncRNA and protein-coding TRGs; down-, up- and mixed regulated TRGs; and early, late and persistent TRGs. (**B**) Proportion of AC and DC features in TRG clusters. AC scores (proportion of TRGs with AC features in TRG cluster) and DC scores (proportion of TRGs with DC features in TRG cluster) classify TRG clusters into AC, DC, AC/DC and NA clusters (background color). Histograms of AC and DC scores are shown. P-values from Shapiro-Wilk test are indicated. (**C**) Genomic size, TRG number and number of TRG neighbors in AC, DC, AC/DC and NA TRG clusters. P-values from Wilcoxon tests are indicated in the plot.

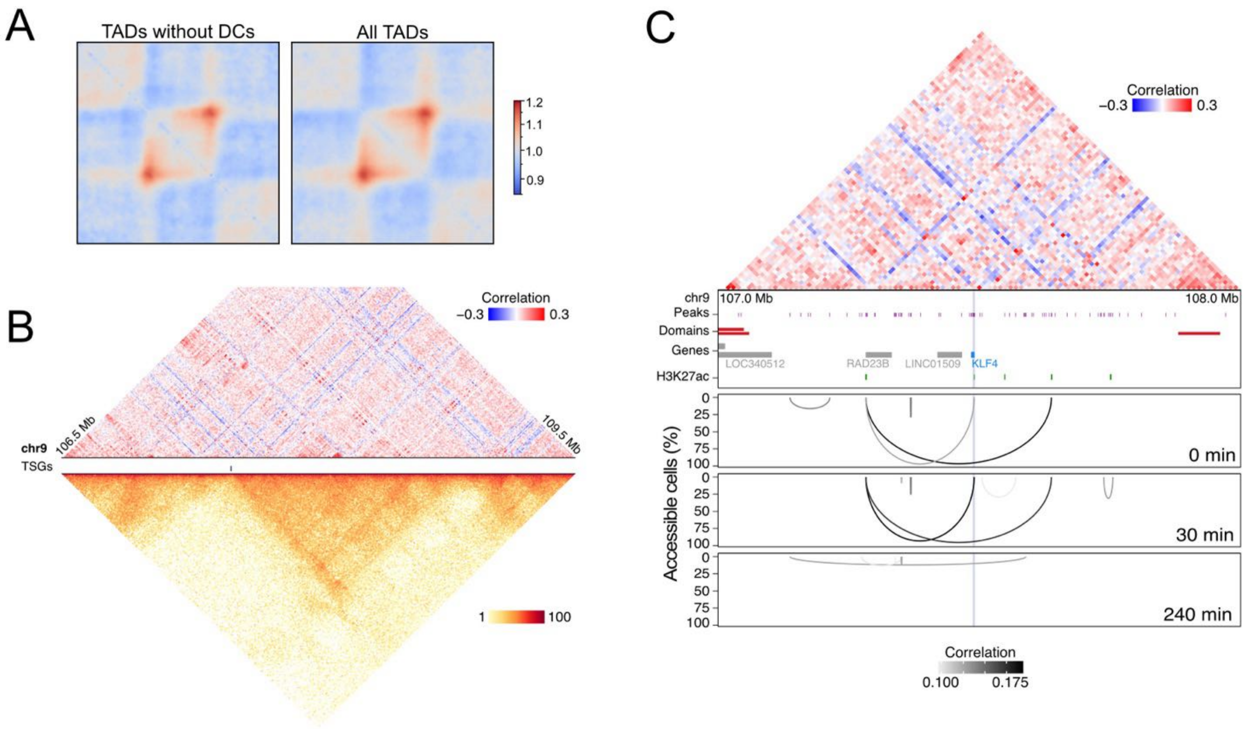


**Supplementary Figure 7. Hi-C chromatin contacts at AC and DC chromatin modules**. (**A**) Chromatin contact pileups of TADs without DCs (left) and all TADs (right) from Hi-C data of unstimulated HUVECs. (**B**) Metacell chromatin co-accessibility and chromatin contacts at the TRG KLF4. Top: co-accessibility map from metacell co-accessibility across unstimulated and TNFα-stimulated HUVECs. The average co-accessibility of replicates is shown. The limits of the color scale are set to -0.3 and 0.3. Middle: genes (gray), TRGs (blue), and 1 kb regions around the TSSs of TRGs (light blue). Bottom: chromatin contact map from Hi-C data of unstimulated HUVECs. The maximum color scale is set to 100. (**C**) Metacell chromatin co-accessibility and ACs from single cell co-accessibility in unstimulated and TNFα-stimulated HUVECs zoomed in at the TRG KLF4. Top: same as in panel B. Middle: 1 kb extended pseudo-bulk ATAC peaks (black); H3K27ac peaks from ChIP-seq of 30 min TNFα-stimulated HUVECs (green); genes (gray), TRGs (blue) and 1 kb regions around the TSSs of TRGs (light blue). Bottom: consensus ACs from single cell co-accessibility of biological replicates. The color and height of loops reflect accessibility correlation and the percent accessible cells of linked peaks.

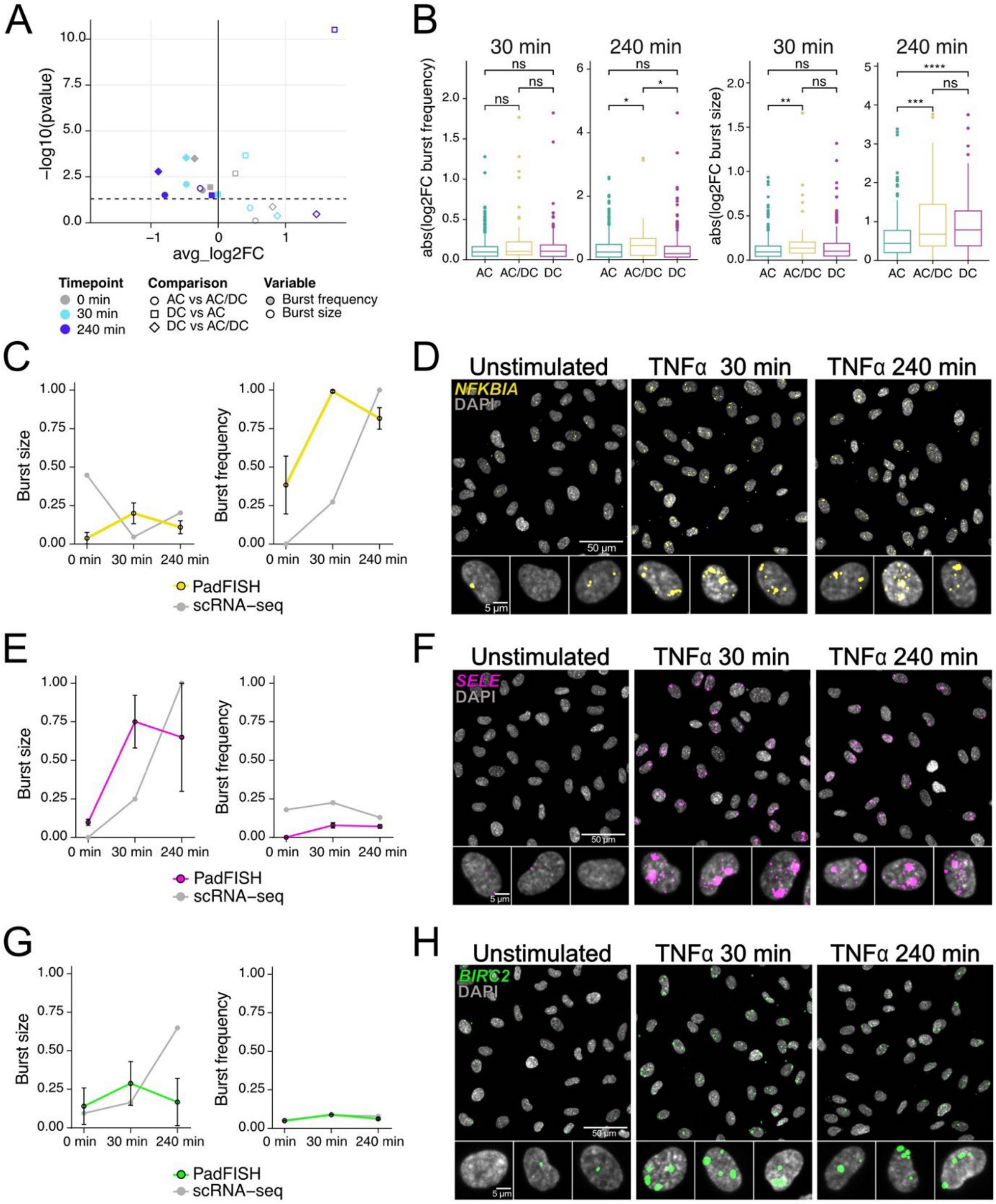


**Supplementary Figure 8. Bursting kinetics of TRGs in AC, DC, and AC/DC chromatin modules.** (**A**) Differential bursting kinetics in TRGs of AC, DC, and AC/DC modules. Average log2FC and p-values from a two-sided Wilcoxon test for differential burst sizes (shape outlines) and frequencies (filled shapes) are shown with p-value = 0.05 indicated by the dashed line. Burst sizes were significantly higher in DC vs. AC, while burst frequencies were significantly higher in AC and AC/DC vs. DC modules. (**B**) Differential bursting kinetics in individual TRGs of AC, DC and AC/DC modules. The absolute log2FC of burst frequencies (left) and sizes (right) per TRG after TNFα treatment are shown. P-values from Wilcoxon test are indicated as p > 0.05, ns; p-value ≤ 0.05, *; p-value ≤ 0.01, **; p-value ≤ 0.001, ***; p-value ≤ 0.0001, ****. Burst size fold changes were significantly higher for TRGs in DC and AC/DC vs. AC, while burst frequency fold changes were significantly higher for TRGs in AC/DC vs AC and DC chromatin modules. (**C**) NFKBIA burst size (left) and frequency (right) from snRNA-seq and padFISH. Error bars represent the standard errors across three biological replicates. (**D**) padFISH images of intronic NFKBIA expression (yellow) and DAPI (gray). Zoom-ins to three exemplary cells per treatment condition are shown. (**E**) Same as panel C for SELE. (**F**) Same as panel D for SELE (magenta). (**G**) Same as panel C for BIRC2. (**H**) Same as panel D for BIRC2 (green).

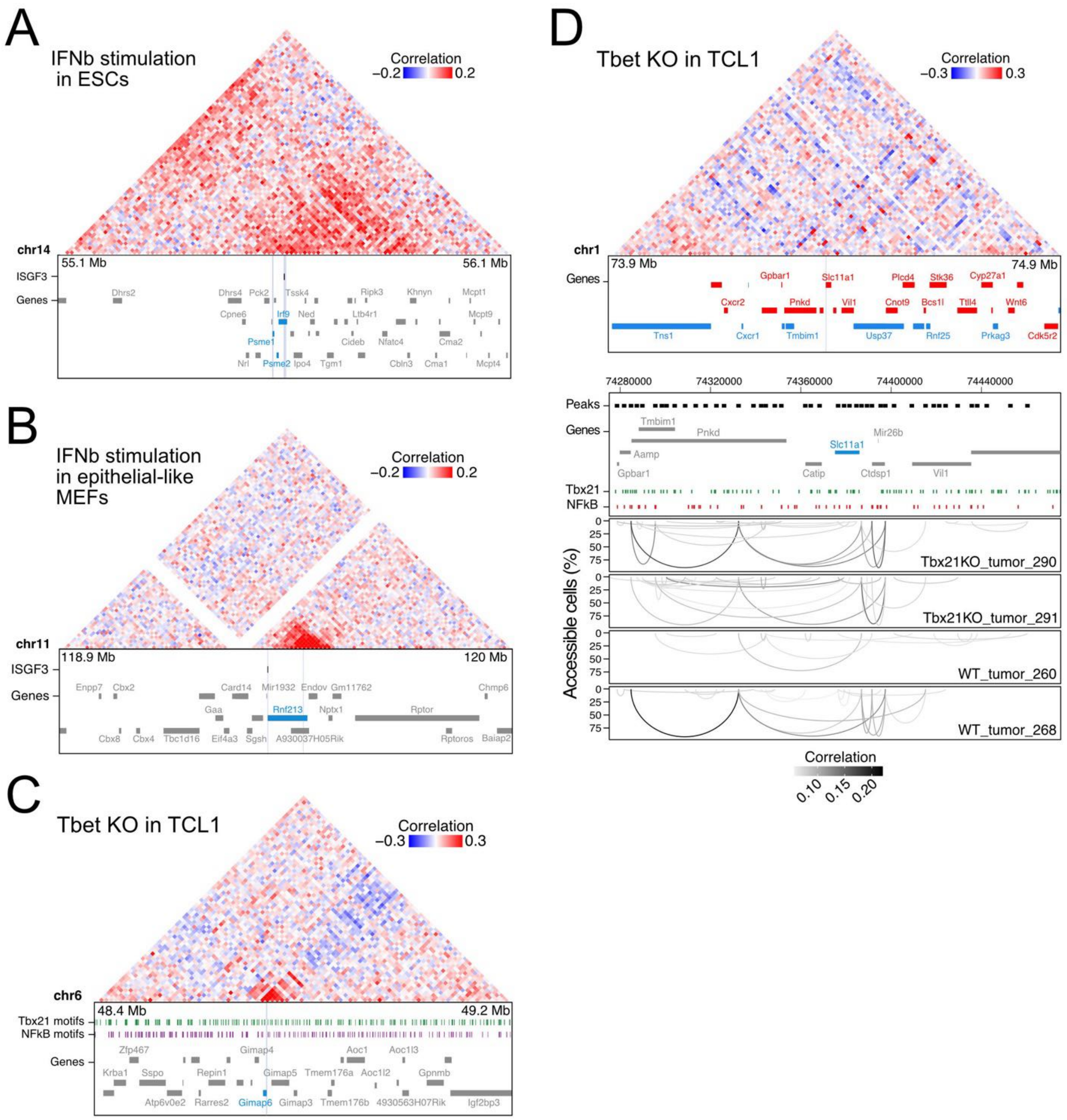


**Supplementary Figure 9. DCs and ACs in mouse cells under perturbation.** (**A**) Metacell chromatin co-accessibility of unstimulated and IFN-stimulated mouse embryonic stem cells at a cluster of interferon stimulated genes (ISG) with Psme1, Psme2 and Irf9. Induced binding of STAT1/2 complexes from ChIP-seq of unstimulated and IFN-stimulated mouse ESCs (black), gene annotation (red, blue) and 1 kb regions around ISG TSSs (gray) are annotated. (**B**) Same as panel A for mesenchymal-like mouse embryonic fibroblasts at ISG Rnf213. (**C**) Metacell chromatin co-accessibility of Tbx21-wt and Tbx21-ko samples of mouse TCL1 model at T-bet dependent gene Gimap6. T-bet binding motifs (black), gene annotation (red, blue) and 1 kb regions around T-bet dependent gene TSSs (gray) are annotated. (**D**) Metacell chromatin co-accessibility and single cell co-accessible links in Tbx21-wt and Tbx21-ko samples of mouse TCL1 model at T-bet dependent gene Slc11a1. Top: co-accessibility map from metacell co-accessibility across Tbx21-wt and Tbx21-ko TCL1 samples. Middle: 2 kb extended pseudo-bulk ATAC peaks (black); Tbet binding motifs (black); gene annotation (red, blue). 1 kb regions around T-bet dependent gene TSSs are marked in gray. Bottom: co-accessible links at T-bet dependent gene promoters from single cell co-accessibility.

#### Supplementary Tables

##### Supplementary Table S1. padFISH of CXCL intronic transcripts

| TNFα (min) | Number/fraction of cells [a] | Rep1 | Rep2 | Rep3 | Mean |
|---|---|---|---|---|---|
| **0** | Total cell population | 465 | 408 | 220 | |
| | Cells with co-localizing transcripts | 11 | 5 | 18 | |
| | **Fraction with co-localization (%)** | **2** | **1** | **8** | **4±4** |
| **30** | Total cell population | 358 | 388 | 271 | |
| | Cells with co-localizing transcripts | 236 | 286 | 163 | |
| | **Fraction with co-localization (%)** | **66** | **74** | **60** | **67±7** |
| **240** | Total cell population | 475 | 393 | 295 | |
| | Cells with co-localizing transcripts | 167 | 122 | 122 | |
| | **Fraction with co-localization (%)** | **35%** | **31%** | **41%** | **36±5** |

Co-expression masks were defined from the co-localization of at least two transcripts. Out of the total cell population analyzed, only those cells that showed a minimum of one and a maximum of two co-expression masks were included in the analysis. The ratio between the two populations (cells with co-localizations vs. total cells) was only 4±4% at 0 min (unstimulated cells) since most cells had no co-localizing spot of two different nascent RNAs. This fraction increased to 67±7% and 36±5% for the 30 min and 240 min time points, respectively.

##### Supplementary Table S2. Inventory of supplementary data sets associated with this manuscript.

| File Name | Description |
|---|---|
| Supplementary Dataset 1: Dataset_01_sc-seq_QC.xlsx | QC parameters of single cell sequencing data. |
| Supplementary Dataset 2: Dataset_02_TRGs.xlsx | TRGs identified in HUVECs after 30 min and 240 min of TNFα treatment with adjusted p-values and log2 fold changes. Information on TRG clustering, factory TRGs and TRG regulation types are provided. |
| Supplementary Dataset 3: Dataset_03_TRGclusters.xlsx | TRG clusters with AC and DC scores and their classification into AC, DC and AC/DC. |
| Supplementary Dataset 4: Dataset_04_ACs.xlsx | Consensus lists of ACs called in at least two replicates. Separate sheets for ACs at different treatment time points are provided. |
| Supplementary Dataset 5: Dataset_05_DCs.xlsx | Called DCs with enriched co-accessibility scores. |
| Supplementary Dataset 6: Dataset_06_PadlockProbes.xlsx | Padlock probes used in the padFISH analysis. |

**Supplementary Table S3. Data analysis packages and software from external sources.**

| Software | Ref | Link | Version |
|---|---|---|---|
| ArchR | 7 | github.com/GreenleafLab/ArchR | 1.0.3 scATAC; 1.0.2 smMultiome |
| Arrowhead | 13 | https://github.com/aidenlab/juicer/wiki/Arrowhead | |
| Bowtie2 | 14 | bowtie-bio.sourceforge.net/bowtie2/index.shtml | 2.3.3 |
| Cell Ranger | 15 | support.10xgenomics.com/single-cell-gene-expression/software/pipelines/latest/what-is-cell-ranger | 7.1.0 |
| Cell Ranger ARC | | https://www.10xgenomics.com/support/software/cell-ranger-arc/latest/getting-started/what-is-cell-ranger-arc | 1.0.1 |
| Cell Ranger ATAC | 16 | support.10xgenomics.com/single-cell-atac/software/pipelines/latest/what-is-cell-ranger-atac | 2.1.0 |
| chromVARmotifs | | https://github.com/GreenleafLab/chromVARmotifs | 0.2.0 |
| cooler | 17 | https://github.com/open2c/cooler | |
| coolpup.py | 18 | https://github.com/open2c/coolpuppy | |
| DESeq2 | 19 | doi.org/10.18129/B9.bioc.DESeq2 | 1.40.2 |
| FIJI | 20 | https://imagej.net/software/fiji/ | 2.14.0 |
| GenomicRanges | 8 | doi.org/10.18129/B9.bioc.GenomicRanges | 1.52.0 |
| gUtils | | doi.org/10.32614/CRAN.package.hutils | 0.2.0 |
| Hic2cool | | https://github.com/4dn-dcic/hic2cool | |
| HiCLift | 21 | https://github.com/XiaoTaoWang/HiCLift | |
| HISAT2 | 22 | https://www.ccb.jhu.edu/software/hisat/index.shtml | |
| Homer | 12 | http://homer.ucsd.edu/homer/ | 4.11 |
| HTSeq | 23 | https://pypi.org/project/HTSeq/ | |
| ilastik | 24 | https://www.ilastik.org | 1.4.0 |
| MACS2 | 25 | https://pypi.org/project/MACS2/ | 2.1.2 |
| Nextflow | 26 | https://nf-co.re/ | 22.10.6 |
| Nf-core rnaseq | | https://nf-co.re/rnaseq/3.14.0/ | 3.9.0 |
| NOISeq | 27 | https://www.bioconductor.org/packages/release/bioc/html/NOISeq.html | 3.19 |
| plotgardener | 9 | https://bioconductor.org/packages/release/bioc/html/plotgardener.html | 1.6.2 |
| poolr | 28 | https://cran.r-project.org/web/packages/poolr/index.html | 1.1-1 |
| Python | | | 3.10.12, TF activity; 3.10.8, snMultiome |
| R | 29 | www.r-project.org | 4.3.1/4.3.2 (Multiome + padFISH) |
| scDblFinder | 30 | doi.org/10.12688/f1000research.73600.2 | 1.14.0, scATAC; 1.16.0, snMultiome |
| SciPy | 31 | scipy.org | 1.11.4 |
| Seurat | 32 | satijalab.org/seurat/ | 4.3.0.1 |
| SpectralTAD | 10 | https://bioconductor.org/packages/release/bioc/html/SpectralTAD.html | 1.16.1 |
| Tobias | 33 | https://github.com/loosolab/TOBIAS | 0.15.1 |

**Supplementary Table S4. Inventory of data sets deposited at external repositories.**

| Data | Repository | Link | Description |
|---|---|---|---|
| scRNA-seq data | GEO, GSE273430 | https://www.ncbi.nlm.nih.gov/geo/GSE273426 | Data for unstimulated and TNFα-stimulated HUVEC samples |
| snATAC-seq data | GEO, GSE273430 | https://www.ncbi.nlm.nih.gov/geo/GSE273428 | Data for unstimulated and TNFα-stimulated HUVEC samples |
| snRNA-seq data | GEO, GSE273430 | https://www.ncbi.nlm.nih.gov/geo/GSE273427 | Data for unstimulated and TNFα-stimulated HUVEC samples |
| snMultiome RNA-/ATAC-seq data | GEO, GSE273430 | https://www.ncbi.nlm.nih.gov/geo/GSE273429 | Data for unstimulated and TNFα-stimulated HUVEC samples |
| RNA-seq (bulk) data | GEO, GSE273430 | https://www.ncbi.nlm.nih.gov/geo/GSE273430 | Data for unstimulated and TNFα-stimulated HUVEC samples |
| H3K27ac ChIP-seq (bulk) data | GEO, GSE273430 | https://www.ncbi.nlm.nih.gov/geo/GSE273430 | Data for unstimulated and TNFα-stimulated HUVEC samples |
| Hi-C (bulk) | GEO, GSE63525 | https://www.ncbi.nlm.nih.gov/geo/GSE63525 | Data for analysis of chromatin contacts in uninduced HUVECs from ref. [13]. |
| snATAC-seq data from mouse ESCs and MEFs | GEO, GSE160764 | https://www.ncbi.nlm.nih.gov/geo/GSE160764 | Data for co-accessibility analysis in mouse ESCs and MEFs from ref. [6]. |
| snATAC-seq data from TCL1 mouse cells | GEO, GSE234226 | https://www.ncbi.nlm.nih.gov/geo/GSE234226 | Data for co-accessibility analysis in TCL1 mouse cells from ref. [11] |
| padFISH | BioImage Archive, S-BIAD1294 | https://www.ebi.ac.uk/biostudies/bioimages/studies/S-BIAD1294 | Fluorescence microscopy source images of the padFISH analysis. |
| Vignette dataset for RWireX (GitHub) | Zenodo | https://doi.org/10.5281/zenodo.13142236 | Vignette dataset for the RWireX software available from Github via the link https://github.com/RippeLab/RWireX |
